# Open modification searching of SARS-CoV-2–human protein interaction data reveals novel viral modification sites

**DOI:** 10.1101/2022.03.10.483652

**Authors:** Charlotte Adams, Kurt Boonen, Kris Laukens, Wout Bittremieux

**Affiliations:** Department of Computer Science, University of Antwerp, 2020 Antwerpen, Belgium; Centre for Proteomics (CFP), University of Antwerp, 2020 Antwerpen, Belgium; Sustainable Health Department, Flemish Institute for Technological Research (VITO), 2400 Mol, Belgium; Skaggs School of Pharmacy and Pharmaceutical Sciences, University of California San Diego, La Jolla, CA 92093, United States of America

**Keywords:** SARS-CoV-2, protein-protein interactions, open modification searching, post-translational modification, phosphorylation, ubiquitination, S-nitrosylation

## Abstract

The outbreak of the SARS-CoV-2 coronavirus, the causative agent of the COVID-19 disease, has led to an ongoing global pandemic since 2019. Mass spectrometry can be used to understand the molecular mechanisms of viral infection by SARS-CoV-2, for example, by determining virus–host protein–protein interactions (PPIs) through which SARS-CoV-2 hijacks its human hosts during infection, and to study the role of post-translational modifications (PTMs). We have reanalyzed public affinity purification mass spectrometry data using open modification searching to investigate the presence of PTMs in the context of the SARS-CoV-2 virus–host PPI network. Based on an over two-fold increase in identified spectra, our detected protein interactions show a high overlap with independent mass spectrometry-based SARS-CoV-2 studies and virus–host interactions for alternative viruses, as well as previously unknown protein interactions. Additionally, we identified several novel modification sites on SARS-CoV-2 proteins that we investigated in relation to their interactions with host proteins. A detailed analysis of relevant modifications, including phosphorylation, ubiquitination, and S-nitrosylation, provides important hypotheses about the functional role of these modifications during viral infection by SARS-CoV-2.

## Introduction

Severe acute respiratory syndrome coronavirus 2 (SARS-CoV-2) is a highly transmissible and pathogenic coronavirus that causes coronavirus disease 2019 (COVID-19). Since late 2019, it has caused an ongoing global pandemic and socioeconomic crisis, with currently over 608 million people infected and over 6.5 million deaths globally (World Health Organization, September 2022) [1]. Despite intensive research by the scientific community, many important questions regarding its molecular mechanisms remain unanswered.

Viruses are opportunistic intracellular pathogens that depend on their interactions with host proteins to ensure their survival and propagation. The study of protein–protein interactions (PPIs) between SARS-CoV-2 and human proteins is important to understand the mechanisms of viral infection by SARS-CoV-2 [2] and develop therapeutic treatments [3]. Proteins and their function can be altered by post-translational modifications (PTMs) to increase the functional diversity of the limited number of proteins that are encoded. Interestingly, PTMs have been characterized on several coronavirus proteins, in spite of the fact that coronaviruses lack enzymes capable of introducing such modifications [4]. Elucidating the roles of PTMs in a mechanistic context is important to understand viral infection, as they are crucial for viral protein function and promote viral replication, assembly, and release. For example, phosphorylation of the SARS-CoV-2 nucleocapsid protein allows its shuttling between cellular compartments [5]. Additionally, insights in viral and host PTM dynamics offers a potential avenue towards the development of antiviral therapies. Removing PTMs that play a role in the enzymatic activity of viral proteins can aid the host in overcoming viral infection. Alternatively, viral proteins can be modified leading to their inactivation and/or proteasomal degradation, for example by attaching ubiquitin [6].

Affinity-purification mass spectrometry (AP-MS)-based proteomics was used to compile the first SARS-CoV-2–human PPI map, revealing interactions with proteins involved in major cellular processes, including DNA replication, RNA processing, and vesicle trafficking to give insights into SARS-CoV-2 infection [2]. Additionally, the interaction map revealed several human proteins that are targeted by existing drugs approved by the US Food and Drug Administration, which may be potential targets for drug repurposing.

The majority of previously conducted studies use standard sequence database searching to process AP-MS data [2, 7–10]. Most search engines require the user to explicitly specify potential protein modifications to be considered during searching. Unfortunately, taking into account multiple modifications simultaneously during spectrum identification is problematic. First, it leads to excessive search times due to the exponential increase in the number of candidate peptides that have to be considered. Second, it produces more random high-scoring matches due to the increase in candidates, leading to fewer identifications at a given false discovery rate (FDR) [11]. Therefore, a standard analysis typically only considers a handful of variable modifications. In the original analysis of the SARS-CoV-2 AP-MS data, only two modifications (N-terminal acetylation and methionine oxidation) were considered, leaving a substantial part of the data unexplored.

We have recently performed a computational reanalysis of the SARS-CoV-2 AP-MS data by Gordon et al. [2] to uncover additional virus–host interactions not reported in the original study, which highlighted further opportunities for drug repurposing [12]. However, post-translational modifications were still not considered. Therefore, in this study we used an increasingly popular approach to overcome these limitations called “open modification searching” (OMS) to gain new insights into virus–host interactions. This innovative approach allows a modified spectrum to match against its unmodified variant by using a very wide precursor mass window. It is thus able to identify peptides carrying any type of modification. Algorithmic advances of the last few years now allow for the fast and accurate use of OMS, enabling an unbiased detection of modifications at an unparalleled scale [13–15]. The OMS solution ANN-SoLo [13, 14] was used to reprocess the SARS-CoV-2 AP-MS data by Gordon et al. [2]. The reanalysis resulted in a more than two-fold increase in identified spectra compared to the originally reported results, allowing more accurate PPI filtering. Additionally, several modified viral peptides were identified. Phosphorylation, ubiquitination, and S-nitrosylation were selected to be investigated in more detail, in view of their putative importance during a viral infection. For each of these PTMs, we detected novel PTM sites on SARS-CoV-2 proteins, revealing potential functional insights.

## Experimental procedures

### Experimental design and statistical rationale

A previously generated AP-MS dataset to study the SARS-CoV-2 virus–host interactome [2] was retrieved from the PRIDE repository (PXD018117) [16]. For the full experimental details, see the original study by Gordon et al. [2]. In brief, affinity purification was performed using 27 SARS-CoV-2 proteins that were individually tagged and expressed in triplicate (biological replicates) in HEK-293T cells. Bead-bound proteins were denatured, reduced, carbamidomethylated, and enzymatically digested using trypsin, and each sample was injected via an Easy-nLC 1200 (Thermo Fisher Scientific) into a Q-Exactive Plus mass spectrometer (Thermo Fisher Scientific). The SARS-CoV-2 proteins that were included are: all mature nonstructural proteins (Nsps), except for Nsp3 and Nsp16; a mutated version of Nsp5 to disable its proteolytic activity (Nsp5_C145A); and all predicted SARS-CoV-2 open reading frames (Orfs), including the spike (S), membrane (M), nucleocapsid (N), and envelope (E) protein. Spectrum identifications were filtered at 1% FDR and PPIs were filtered using a SAINTexpress Bayesian false-discovery rate (BFDR) ≤ 0.05, an average spectral count ≥ 2, and a MiST score ≥ 0.7 (see further).

### Spectrum identification using open modification searching

First, the downloaded raw files were converted to MGF files using ThermoRawFileParser (version 1.2.3) [17]. Next, OMS was performed using the ANN-SoLo spectral library search engine (version 0.2.4) [13, 14]. A combined human–SARS-CoV-2 spectral library was used for searching. The MassIVE-KB library (version 2018/06/15) was used as human spectral library. This is a comprehensive human HCD spectral library containing 2,154,269 unique precursors corresponding to 1,114,503 unique peptides, derived from publicly available mass spectrometry data in the MassIVE repository [18]. SARS-CoV-2 spectra were simulated by generating all possible tryptic peptide sequences from the SARS-CoV-2 protein sequences downloaded from UniProt (version 2020/03/05) using Pyteomics (version 4.3.2) [19] and predicting the corresponding spectra using Prosit (version prosit_intensity_2020_hcd; collision energy 33 as determined by Prosit collision energy calibration) [20]. A simulated spectral library for the green fluorescent protein was generated in a similar fashion. A final spectral library was compiled by merging all spectra using SpectraST (version 5.0) [21] and adding decoy spectra in a 1:1 ratio using the shuffle-and-reposition method [22]. ANN-SoLo was configured to use a 20 ppm precursor mass tolerance during the first step of its cascade search and a 500 Da precursor mass tolerance during its open search. Other search settings were to filter peaks below 101 *m*/*z*, above 1500 *m*/*z*, and in a 0.5 *m*/*z* window around the precursor mass; a 0.02 *m*/*z* fragment mass tolerance; and a bin size of 0.05 *m*/*z*. The remaining settings were kept at their default values. Peptide-spectrum matches (PSMs) were filtered at 1% FDR using ANN-SoLo’s built-in subgroup FDR procedure (Supplementary Table 1).

Additionally, OMS was performed using MSFragger (version 3.5) [15] and FragPipe (version 18.0) against a concatenated FASTA file containing human protein sequences (Uniprot reviewed sequences downloaded on 2020/02/28) [23], the SARS-CoV-2 protein sequences (version 2020/03/05), and the green fluorescent protein sequence. An equal number of decoy protein sequences were generated using FragPipe. The MSFragger search settings included a precursor mass tolerance between −150 Da and 500 Da, a fragment mass tolerance of 0.02 Da, and trypsin cleavage with up to two missed cleavages. Cysteine carbamidomethylation was used as a fixed modification, and oxidation of methionine and N-terminal acetylation were used as variable modifications. Other search settings were kept at their default values. PSMs were processed using PeptideProphet (version 4.4.0) [24] with the FragPipe default settings for open searches and filtered at 1% FDR.

### Protein inference and protein-protein interaction filtering

Protein inference was performed using the Protein Inference Algorithms (PIA) tool (version 1.3.13) [25] based on Occam’s razor. Only proteins with minimum two unique peptides were retained, while other settings were kept at their default values (Supplementary Table 2). Combined scoring of interacting proteins using SAINTexpress (version 3.6.3) [26] and MiST (https://modbase.compbio.ucsf.edu/mist/, version main.e2da2b0) [27] was used to filter high-confidence PPIs (Supplementary Table 3). Scoring thresholds were a SAINTexpress Bayesian false-discovery rate (BFDR) ≤ 0.05, an average spectral count ≥ 2, and a MiST score ≥ 0.7 (Supplementary Table 4).

To validate the filtered PPIs, a list of SARS-CoV-2–human interactions reported in seven alternative MS-based SARS-CoV-2 virus–host interactome studies [7–10, 28–30] were obtained from the BioGRID repository [31]. Additionally, PPIs from a previous reanalysis of the original AP-MS study were included [12]. Human interaction partners were queried in the VirHostNet database (version 3.0) [32] to investigate whether these proteins are also targeted by other viruses.

### Gene ontology enrichment analysis

Gene Ontology (GO) enrichment analysis was performed for the human proteins that interact with each viral protein, using the enrichGO function of the clusterProfiler package (version 4.0.5) in R. Significant GO terms corresponding to the biological process category (1% Benjamini-Hochberg false discovery rate) were extracted and further refined to select non-redundant terms using the rrvgo package (version 1.4.4) with default parameters [33].

### Investigation of post-translational modifications

The precursor mass differences observed in the ANN-SoLo results were referenced against the Unimod database [34] to determine the modifications that were present. PSMs that included specific PTMs (phosphorylation, ubiquitination, and S-nitrosylation) were manually investigated in more detail to disambiguate between alternative PTM assignments with near-identical mass and determine the modification site by visual inspection using the spectrum_utils Python package (version 0.3.3) [35].

Identified PTM sites were verified using PTM prediction tools and through literature study. Phosphorylation results were compared to three independent SARS-CoV-2 phosphoproteomic studies [36–38]. Phosphorylation site prediction was performed using NetPhos (version 3.1) [39, 40]. Additionally, GPS (version 5.0) [41] was used to predict kinase-specific phosphorylation sites (with a “high” threshold) for each of the SARS-CoV-2 proteins that were found to interact with known human kinases, retrieved from the KinHub database [42] (Supplementary Table 5–6). Ubiquitination results were compared to two independent SARS-CoV-2 ubiquitination studies [7, 43] and ubiquitination sites predicted by BDM-PUB (version 1.0) [44] (Supplementary Table 7). To our knowledge, to date there have been no S-nitrosylation sites reported on SARS-CoV-2. S-nitrosylation site prediction was performed with iSNO-PseAAC (version 1.0) [45], to validate observed S-nitrosylation sites (Supplementary Table 8–9).

## Results

### Increased spectrum identification rate using open modification searching boosts protein–protein interaction confidence

Open modification searching using ANN-SoLo succeeded in identifying 830,743 of 2,503,010 total MS/MS spectra at 1% FDR (33% identification rate). This represents a 214% increase in identified spectra compared to the originally reported results [2] obtained by standard searching using MaxQuant [46] (Figure 1A). Notably, 402,586 PSMs correspond to modified peptides with non-zero precursor mass differences from open modification searching (Figure 1C, Supplementary Table 10). Besides the increase in identified spectra, an important advantage of open modification searching is its ability to identify any type of PTM in an unbiased fashion, without the need to explicitly specify a limited number of variable modifications. This makes it possible to explore the general presence of PTMs in the context of the SARS-CoV-2 virus–host interactome. Frequently observed PTMs include modifications that were likely artificially introduced during sample processing [47], such as oxidation, dioxidation, and acetylation. Such ubiquitous modifications are typically included as variable modifications during standard searching. The OMS results also include unique observations of biologically relevant modifications at lower abundances, such as phosphorylation, ubiquitination, and S-nitrosylation.

**Figure 1.**
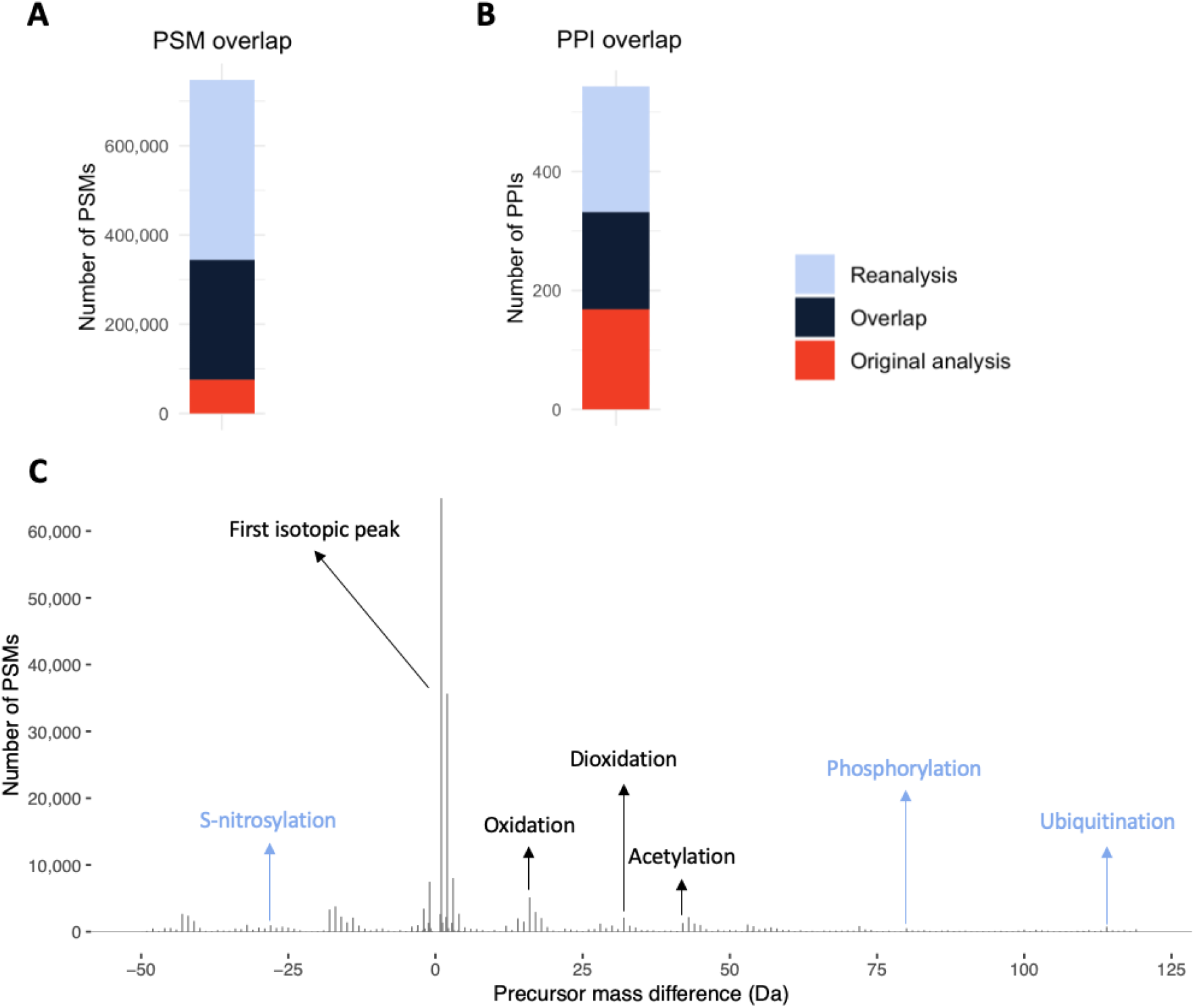
**(A)** Comparison in identification performance between the originally reported results using standard searching [2] and open modification searching. Open modification searching was able to identify more than twice as many spectra, corresponding to the identification of a large number of modified PSMs. **(B)** Comparison of the number of filtered PPIs. Half of the previously reported PPIs were reproduced [2], while 211 new PPIs were determined based on the extended identification results obtained using open modification searching. **(C)** Modifications can be derived from the precursor mass differences observed by open modification searching. Some of the most frequent delta masses are annotated with their likely modifications, sourced from Unimod [34], with modifications of artificial origin in black and relevant biological modifications in blue. The full list of observed precursor mass differences and their likely modifications is available in Supplementary Table 10.

Similar beneficial results can also be achieved with alternative OMS tools, such as MSFragger [15]. Performing an open search using MSFragger instead of ANN-SoLo succeeded in identifying 574,394 of 2,503,010 total MS/MS spectra at 1% FDR (23% identification rate). This represents a 148% increase in identified spectra compared to the originally reported results [2] (Supplementary Figure 1A). Furthermore, there is a strong correspondence in identification results between ANN-SoLo and MSFragger (Supplementary Figure 1B–C), which indicates the robustness of open modification searching, irrespective of the search engine that is employed. For simplicity of the downstream analyses and because ANN-SoLo was more sensitive than MSFragger, the ANN-SoLo results were used to investigate protein interactions during viral infection by SARS-CoV-2.

PPI filtering is crucial to separate true interactors from non-specific binders and contaminants. A combination of SAINTexpress and MiST filtering of the open modification searching results produced 375 high-confidence PPIs (Supplementary Table 4). These results contain 164 PPIs that overlap with the previously reported results [2] and 211 novel PPIs (Supplementary Figure 2). Notably, a previous reanalysis of these AP-MS data using alternative bioinformatics tools reported a similar overlap with the original PPI results [12]. The difference in detected PPIs is partly due to the difference in spectrum identification and PPI filtering strategies. In the original analysis a two-step filtering strategy was used. In the first step, the PPIs were filtered by a SAINTexpress BFDR ≤ 0.05, an average spectral count ≥ 2, and a MiST score ≥ 0.7. All proteins that fulfilled the first filtering step were searched in the CORUM [48] database of known protein complexes, and information was extracted about the stable protein complexes that they participated in. In the second step, all the proteins that formed known complexes with interactors identified in the first step were subjected to a filtering step with a lower stringency (MiST score ≥ 0.6) [2]. In contrast, in the reanalysis only a single filtering step was performed using a SAINTexpress BFDR ≤ 0.05, an average spectral count ≥ 2, and a MiST score ≥ 0.7.

When compared to the unfiltered PPI data of the original analysis [2], 199 of the 211 novel PPIs were previously detected as well but failed the original PPI filtering thresholds (Supplementary Figure 3). Most discarded PPIs (~88%) did not pass the minimum threshold for the MiST score, which is a linear combination of the prey abundance, the prey reproducibility across repeated runs, and the specificity of the prey relative to other baits [49]. About 28% of the discarded PPIs did not pass the SAINTexpress BFDR filter, which is based on the abundance of the preys and control proteins [26]. The difference in filtered PPIs in the reanalysis is driven by the increased number of PSMs per identified protein from the open modification results, which resulted in a more reliable identification of both background proteins and potential interaction partners. This influences the PPI filtering and results in a more robust identification of true interactors.

The PPIs were validated against seven alternative MS-based SARS-CoV-2 virus–host interactome studies [7–10, 28–30] and the results from a previous reanalysis [12]. The largest overlap was found with the previous reanalysis and the original analysis, as these studies make use of the same experimental data and only differ in the computational tools that were employed (Figure 2A). Although most independent studies contributed unique PPIs, there is a strong correspondence in the superset of detected PPIs across all independent studies, with 317 of the 375 PPIs that were detected in this study previously reported in one or more of the other studies (Figure 2A, Supplementary Figure 2). Additionally, investigating the overlap with the VirHostNet database [32] shows that 328 of the human interaction partners are also targeted by other viruses (Figure 2B). These results are significantly enriched in the VirHostNet database (Fisher’s exact test, p-value ≪ 0.001), which is expected as different viruses have similar strategies to hijack the host, for example, through functional conservation or viral recombination [32].

**Figure 2.**
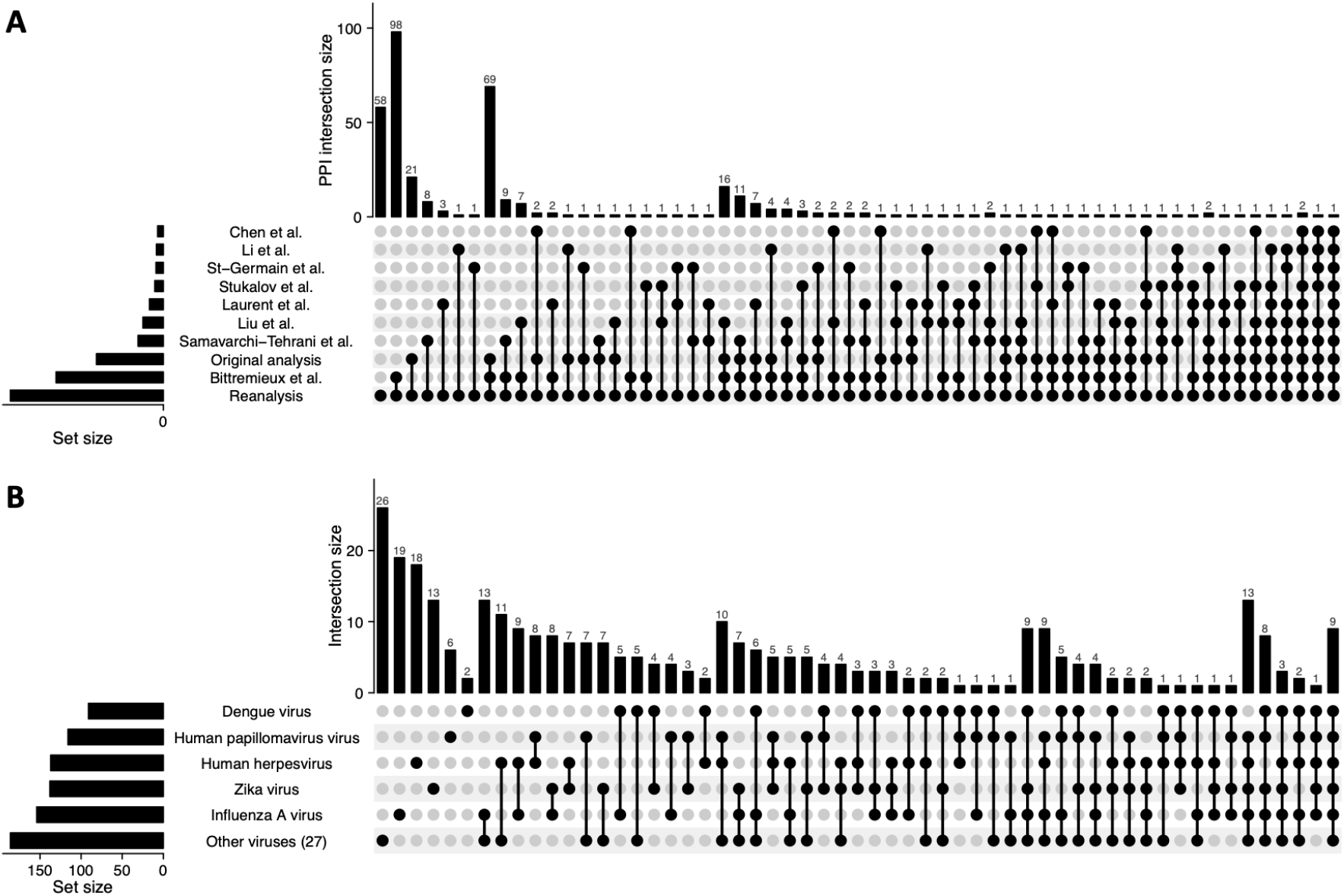
**(A)** Upset plot showing the overlap with seven alternative MS-based SARS-CoV-2 virus–host interactome studies, the original analysis, and a previous reanalysis of the AP-MS data performed by Bittremieux et al. [12]. We observe the largest overlap in PPIs with the previous reanalysis and the original analysis, as all are based on the same AP-MS data. However, several novel detected PPIs have been observed in independent SARS-CoV-2 interactome studies as well, boosting the confidence in these interactions. **(B)** Upset plot showing the overlap of the detected human interaction partners with targets reported for different viruses in the VirHostNet database (excluding VirHostNet data for MERS-CoV, SARS-CoV-1, and SARS-CoV-2).

GO enrichment analysis based on the human interacting proteins for each SARS-CoV-2 protein indicates how viral infection might hijack major cellular processes, including metabolic processes involving non-coding RNA (ncRNA; Nsp8) and glycoproteins (Orf8), and RNA export from the nucleus (Orf6) (Figure 3). Both microRNA (miRNA) and long non-coding RNA (lncRNA) are ncRNA that can regulate gene expression and exhibit different expression profiles in COVID-19 patients compared to healthy people [50, 51]. Additionally, they have recently been found to act as viral modulators, regulating viral infection and host defense [52, 53]. Glycoprotein metabolic processes are expected to be relevant to Orf8, as induction of proinflammatory cytokine production by secreted Orf8 is glycosylation dependent [54]. RNA export from the nucleus is also relevant, as SARS-CoV-2-infected cells have an increased level of nuclear mRNA accumulation. Confirming our detected interactions between Orf6 and the mRNA export factors Rae1 and Nup98, a recent study found that Orf6 uses this process to trap host mRNA [55]. In addition to these major cellular processes, other biological processes relevant during viral infection were found, including viral transcription and viral gene expression (Supplementary Figure 4).

**Figure 3.**
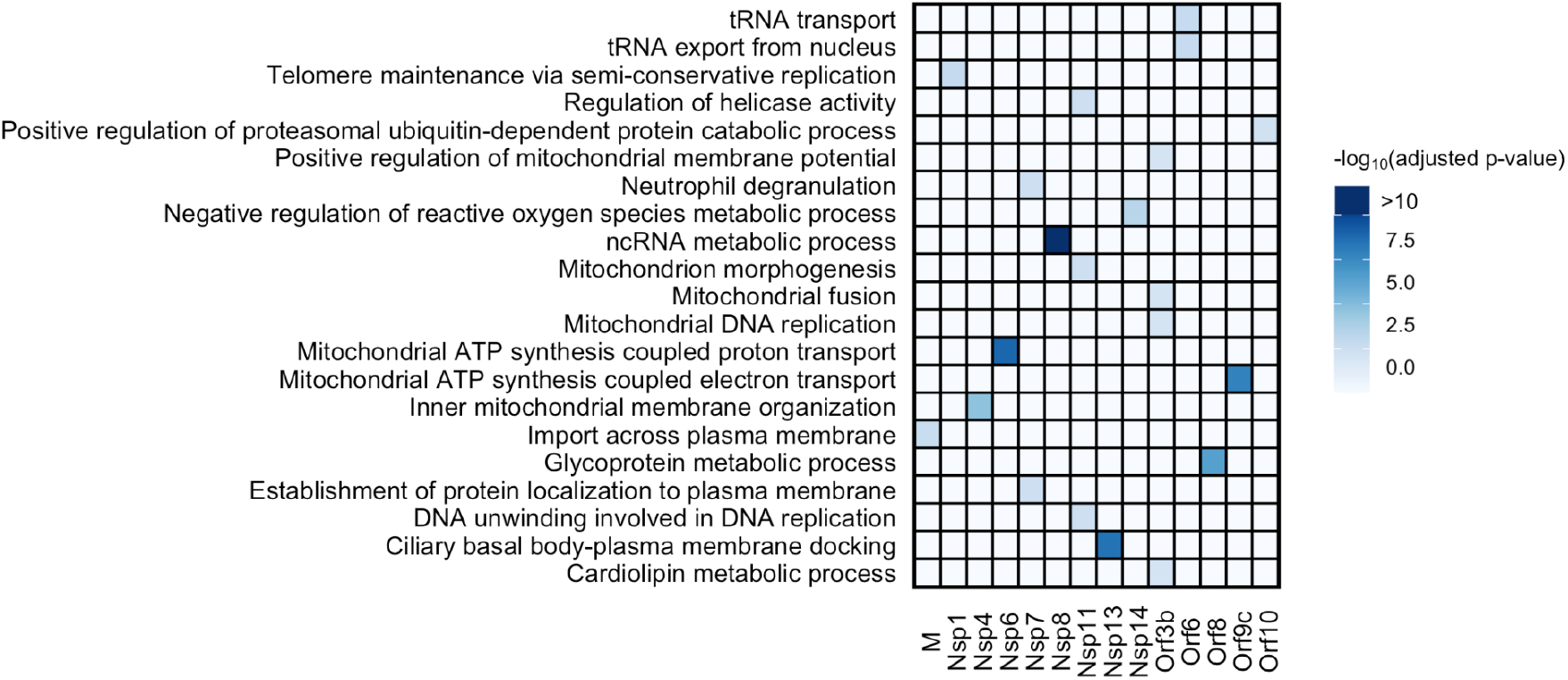
Heatmap showing the top GO terms from the GO enrichment analysis for each of the SARS-CoV-2 proteins.

### Interaction between human kinases and phosphorylated viral proteins

Phosphorylation is a common PTM that impacts many basic cellular processes [56]. It usually results in a functional change of the target protein, interfering with its enzymatic activity, cellular location, and/or association with other proteins. Several studies have indicated that also the function of viral proteins can be affected by their phosphorylation status. For example, association of the SARS-CoV-2 nucleocapsid protein with the 14-3-3 host proteins depends on its phosphorylation status [57] and phosphorylation of the SARS-CoV-1 nucleocapsid protein has been proven to be important for the regulation of the viral life cycle [58].

Twenty phosphorylated viral peptides were identified using open modification searching, and phosphosites were localized to specific residues using visual inspection where possible (Supplementary Table 5, Supplementary Figure 5–24). Notably, despite the fact that no phosphopeptide enrichment protocol was used during the original AP-MS study, we identified several phosphorylation sites that match results from specialized phosphoproteomics studies performed on SARS-CoV-2 infected cells [36–38] (Figure 4A).

**Figure 4.**
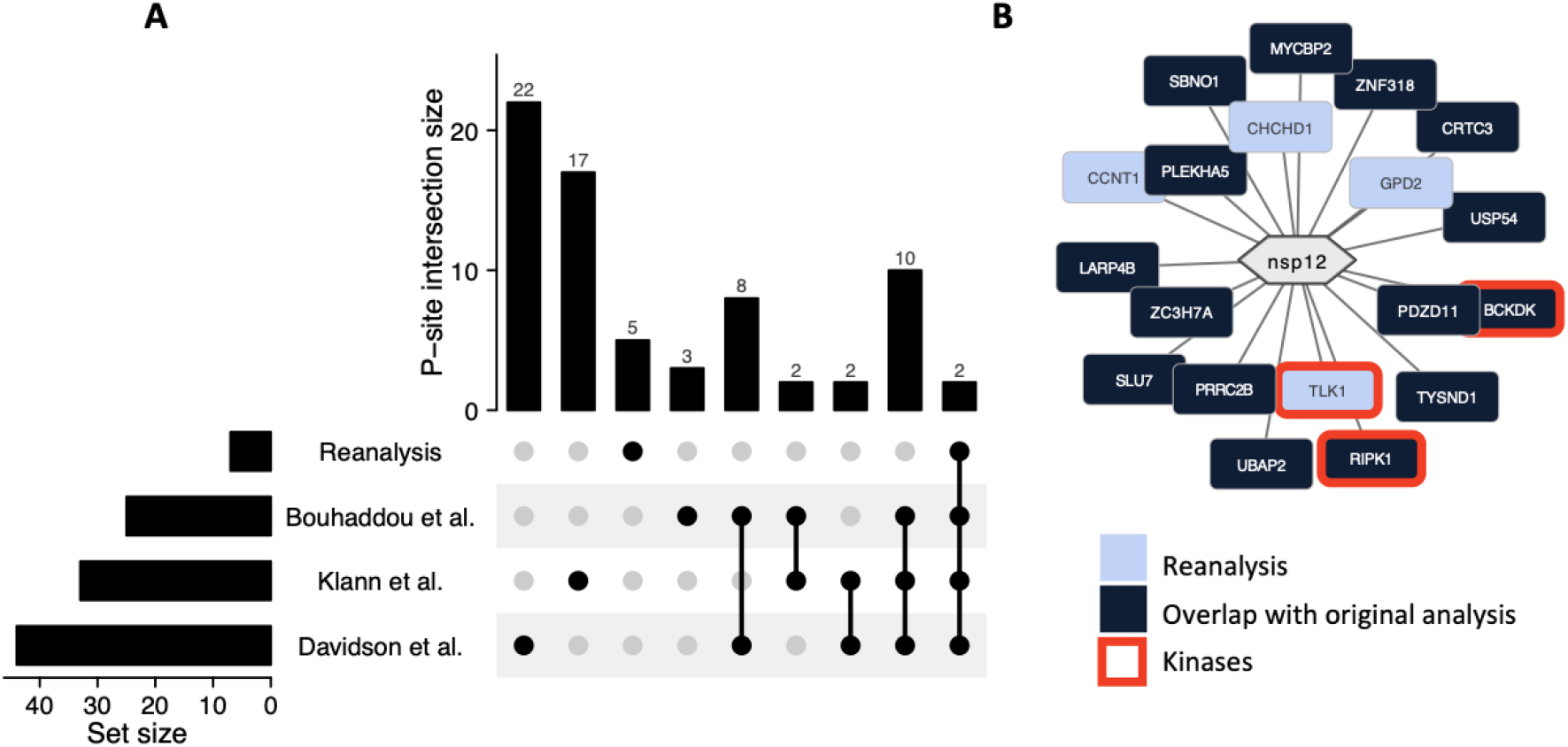
**(A)** Upset plot showing the overlap in phosphorylation sites between independent phosphoproteomic studies and the sites found in the current AP-MS reanalysis. **(B)** The PPI network of Nsp12 with highlighted kinases. Although RIPK1 and BCKDK were found in the original analysis, to our knowledge TLK1 has not been identified as an interaction partner of Nsp12 yet. After kinase-specific phosphorylation site prediction, we found that one of the observed phosphorylation sites (T21) is predicted to be phosphorylated by TLK1.

Phosphorylation status is determined by the interplay between kinases and phosphatases. Kinases catalyze the attachment of phosphate groups to target proteins and phosphatases remove phosphate groups from target proteins [59]. SARS-CoV-2 does not encode any kinases or phosphatases, and thus relies on host proteins for its phosphorylation status. Several kinases and phosphatases have been identified as potential drug targets and a few kinase inhibitors are currently being used to treat COVID-19. For example, baricitinib is recommended by the World Health Organization (WHO) for patients with severe or critical COVID-19. It is a JAK inhibitor that suppresses overstimulation of the immune system by preventing phosphorylation of key proteins involved in the signal transduction that leads to immune activation and inflammation [60], and it is able to prevent SARS-CoV-2 from entering the cell by inhibiting clathrin-mediated endocytosis [61].

We detected multiple interactions between SARS-CoV-2 proteins and host kinases and phosphatases (Supplementary Tables 11–12), including RIPK1 (Nsp12) and TBK1 (Nsp13), both of which have previously been linked to SARS-CoV-2. Active phosphorylated RIPK1 was found in epithelial cell samples from COVID-19 patients [62], and a recent study discovered that Nsp12 promotes activation of RIPK1, which in turn enhances viral replication by stimulating the expression of viral receptors, such as ACE2 and EGFR [63]. Furthermore, Nsp13 is found to limit the activation of TBK1 by directly binding to TBK1 [64]. Additionally, we uncovered previously unknown interactions with host kinases, such as the interaction between Nsp12 and TLK1 (Figure 4B). This interaction is particularly interesting as kinase-specific phosphorylation site prediction revealed that an observed phosphorylation site on Nsp12 (T21) is most likely caused by TLK1.

### Ubiquitination of SARS-CoV-2 proteins

Ubiquitination is a common PTM that can affect the localization, stability, and function of proteins [65]. It regulates a variety of cellular processes, including protein degradation, protein trafficking, transcription, cell-cycle control, and cell signaling [66]. Depending on the cellular context, ubiquitin attachment may either promote or inhibit the viral life cycle. Viruses have developed means to exploit protein ubiquitination by enhancing or inhibiting ubiquitination of specific substrates depending on their needs. They can redirect protein degradation towards proteins with antiviral activity [67] or use ubiquitination to regulate viral proteins. For example, transcriptional function of the human immunodeficiency virus type-1 (HIV-1) Tat protein is increased by the addition of a single ubiquitin molecule [68]. More recently, it has been discovered that the inhibition of IFN-α signaling by SARS-CoV-2 Orf7a depends on the ubiquitination of K119 [69].

Ten ubiquitinated viral peptides were identified, and, where possible, ubiquitination sites were localized to specific residues using visual inspection (Supplementary Table 7, Supplementary Figure 25–34). Notably, all ubiquitination sites that could be confidently localized have been previously reported in independent studies [7, 43] (Figure 5A) or could be predicted. Interestingly, although ubiquitination of K16 on Orf9c has not been reported in SARS-CoV-2 yet, this site has been observed to be ubiquitinated in SARS-CoV-1 [7].

**Figure 5.**
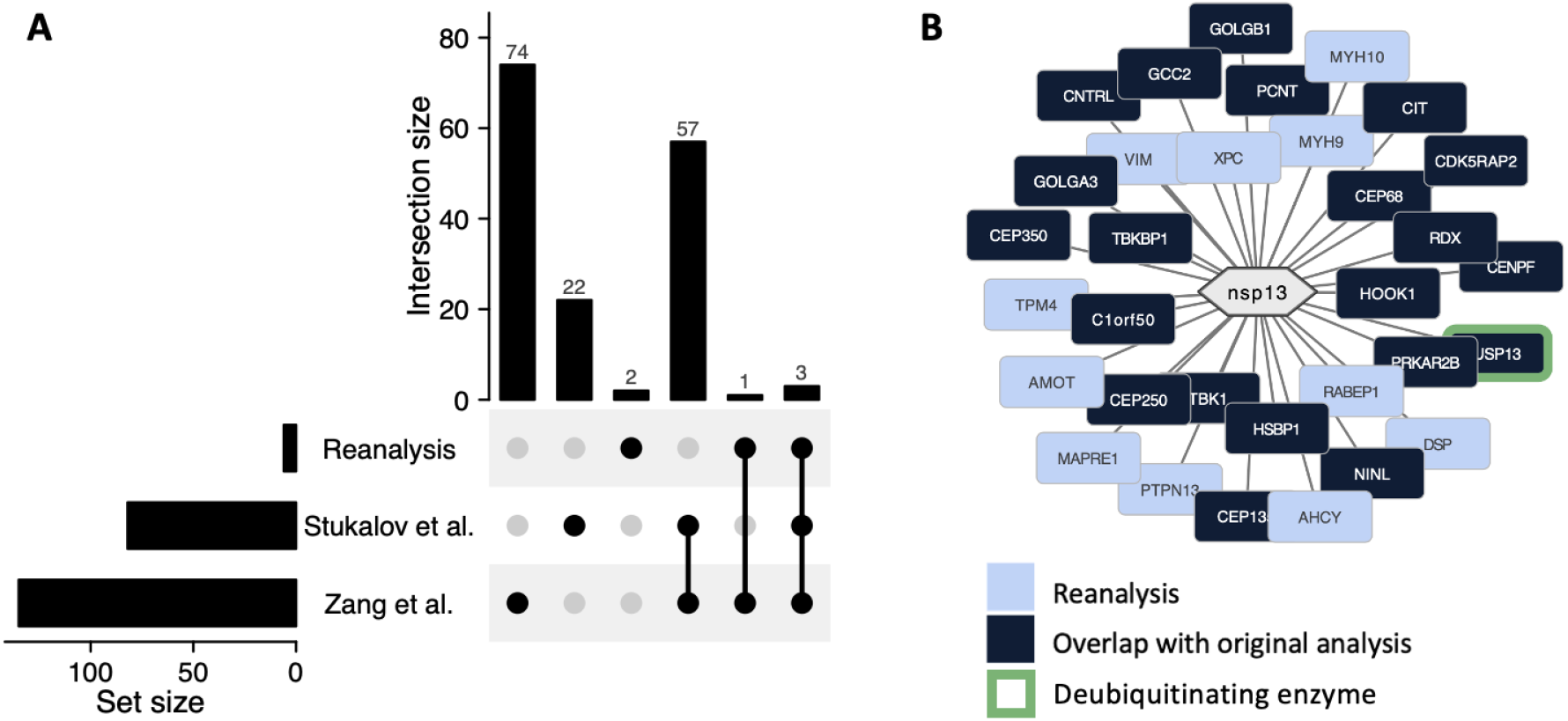
**(A)** Upset plot showing the overlap in ubiquitination sites between independent ubiquitination studies and the sites found in the current AP-MS reanalysis. The two novel ubiquitination sites were verified through *in silico* prediction. K16 on Orf9c has not been reported for SARS-CoV-2 yet, but was previously observed in SARS-CoV-1. **(B)** The PPI network of Nsp13 with the deubiquitinating enzyme USP13 highlighted.

The final step of ubiquitination is performed by a ubiquitin (E3) ligase. This enzyme is particularly important as it determines to which substrate protein the ubiquitin is attached. A ubiquitin can be removed by deubiquitinating enzymes (DUBs). Both host E3 ligases and DUBs were found to interact with SARS-CoV-2 proteins (Supplementary Tables 13–14), suggesting that SARS-CoV-2 might hijack the host ubiquitination system. For example, Nsp13 was found to interact with the DUB USP13 (Figure 5B). According to a recent study, Nsp13 likely hijacks USP13 to prevent itself from degradation. Both knockdown of USP13 and its inhibition through spautin-1 resulted in decreased levels of Nsp13, suggesting that USP13 deubiquitinates and consequently stabilizes Nsp13 [70].

### S-nitrosylation of SARS-CoV-2 proteins

A promising compound currently undergoing clinical trials for COVID-19 is nitric oxide (NO) [71]. Besides its role as an important vasodilator, to prevent blood clot formation, it functions as a vital immune mediator, exerting broad spectrum antiviral effects [72]. Nitric oxide potentially prevents infection by SARS-CoV-2, as it was suggested for SARS-CoV [73]. The surprisingly low prevalence of smokers among hospitalized COVID-19 patients [74] could be attributed to the intermittent burst of high nitric oxide concentration in cigarette smoke [75]. Also nitrate-rich nutrition, exercise, and breathing through your nose is hypothesized to prevent a SARS-CoV-2 infection, as they all increase the nitric oxide concentration [73, 76].

The general antiviral mechanism appears to be the NO-mediated S-nitrosylation of viral and host proteins [77]. S-nitrosylation is the reversible, covalent attachment of nitric oxide to the thiol side chain of cysteine. It is one of the most important and universal PTMs, and it can act as a global regulator of protein function akin to phosphorylation and ubiquitination [78]. Interestingly, reactive cysteine residues are present in many viral and host proteins, representing possible targets for nitric oxide [77]. Results of an *in vitro* study suggest that S-nitrosylation of the SARS-CoV-2 3CL protease (Nsp5) can directly inhibit its protease activity and reduce viral replication [79].

To the best of our knowledge, there are currently no reported S-nitrosylation sites on SARS-CoV-2 proteins. Because cysteine residues are typically considered using a fixed carbamidomethylation modification (+57.021464 Da) introduced during sample processing by alkylation with iodoacetamide [47], Prosit [20]—which was used to simulate SARS-CoV-2 spectra—always considers cysteine residues to be carbamidomethylated. However, the presence of a prior modification can block reduction and alkylation of cysteine residues [80]. Therefore, rather than identifying S-nitrosylation directly based on its mass difference of +28.990164 Da, this modification is represented by a mass difference of −28.031300 Da (28.990164 Da - 57.021464) in the open modification searching results. Because the corresponding S-nitrosylation mass difference is identical to the mass difference observed from the valine to alanine and methionine to cysteine amino acid substitutions, careful visual inspection to localize the modification to specific amino acid residues was performed to confirm three S-nitrosylation sites (Supplementary Table 8, Supplementary Figure 35–52).

Interestingly, one of these S-nitrosylation sites (C45) is located on the main protease Nsp5, specifically, near the catalytic site. Nsp5 is one of two cysteine proteases necessary for viral replication and assembly, and its protease activity is suggested to be directly inhibited by S-nitrosylation [79]. A recent study found that C45 is a hyper-reactive cysteine with a higher nucleophilicity than C145—the catalytic cysteine of Nsp5—and identified C45 as an attractive binding site for the development of a covalent inhibitor [81].

## Discussion

Mass spectrometry research can help to provide insights into the etiology of SARS-CoV-2 infection and identify potential therapeutic targets by investigating the roles of viral and host proteins during infection, their protein interactions, and post-translational modifications (PTMs) [12]. Additionally, it can be used to develop diagnostic strategies, such as through efforts of the CoV-MS consortium [82]—a partnership by multiple academic and industrial groups to increase applicability, accessibility, sensitivity, and robustness of a mass spectrometry-based diagnostic test that detects proteolytically digested SARS-CoV-2 proteins.

Here we have used open modification searching to reprocess SARS-CoV-2–human AP-MS data to investigate post-translational modifications in the context of protein–protein interactions between SARS-CoV-2 and its human host. Studying these two aspects in tandem is especially relevant to understand the mechanisms of viral infection, because although PTMs are essential for viral replication, coronaviruses lack the enzymes to introduce these themselves. Whereas most SARS-CoV-2 studies so far have only considered a limited number of modifications commonly introduced during sample processing, our open modification searching strategy enabled the unbiased investigation of any PTM. This increased the number of identified spectra by more than two-fold, corresponding to newly identified modified peptides, and enabled us to put these PTMs in the context of the virus–host PPI network.

We discovered several combinations of relevant protein interactions and PTMs that hint at functional roles during viral infection. Specifically, we investigated phosphorylation, ubiquitination, and S-nitrosylation in more detail. These are reversible PTMs that are essential for a variety of cellular processes and could be of importance during viral infection. We found interactions between phosphorylated SARS-CoV-2 proteins and host kinases, between ubiquitinated SARS-CoV-2 proteins and host E3 ligases, and novel S-nitrosylation sites on SARS-CoV-2 proteins. Notably, even though no specialized modification enrichment was performed, we were able to confidently detect and localize multiple PTM sites that have been previously reported in independent enrichment studies, as well as novel PTM sites on SARS-CoV-2 proteins.

An important consideration in this study is the overlap in PPI results with the original study [2], which used standard searching instead of open modification searching. Although the vast majority of original spectrum identifications (79%) could be replicated using open modification searching, the overlap in PPIs was smaller (49%). Notably, another recent reanalysis of these data using alternative bioinformatics software tools showed a similarly limited overlap with the original PPI results [12]. Besides using stringent FDR control and other PPI filtering settings during all data processing steps, we validated the detected protein interactions through comparison with alternative MS-based SARS-CoV-2 studies and general virus–host interactions in the VirHostNet database. This showed a highly significant overlap between our PPI results and these independent sources, reinforcing the validity of the newly detected interactions.

The partial mismatch in PPI results can be largely explained by more complete identification of both background proteins and potential interaction partners using OMS. As such, the current study highlights the dependence of PPI filtering on the preceding protein identification results. First, open modification searching can be used to obtain high-quality identification data from which comprehensive PPI results can be obtained. Second, PPI filtering algorithms have to be able to robustly deal with uncertainty and missingness in the identification results when determining true protein interactions. Because OMS increased both the robustness of the PPI filtering results and enabled us to put PTM information in context of the protein interaction network, we suggest that this strategy is ideally suited for the analysis of MS-based PPI data.

In conclusion, we have used open modification searching to reanalyze open SARS-CoV-2 virus–host protein interaction data. The presented results enrich our knowledge of viral infection by SARS-CoV-2 by putting post-translational modifications in the context of the virus–host PPI network, which provides important hypotheses on the functional roles of PTMs during SARS-CoV-2 infection.

## Data availability

All reanalysis results have been deposited to the ProteomeXchange Consortium [83] via the MassIVE repository with data set identifiers RMSV000000359.1 and RMSV000000309.4. Annotated spectra can be inspected in MS-Viewer (https://msviewer.ucsf.edu/prospector/cgi-bin/msform.cgi?form=msviewer) using the key mcorgdznfy. All data processing code is freely available on GitHub as open source under the Apache 2.0 license at https://github.com/adamscharlotte/SARS-Cov-2-analysis.

## Acknowledgements

This research was supported by the University of Antwerp BOF DOCPRO, FWO G070722N, and IBOF VLIR MIMICRY. KL acknowledges support of the University of Antwerp Core Facility BIOMINA. The computational resources and services used in this work were provided by the HPC core facility CalcUA of the University of Antwerp, and VSC (Flemish Supercomputer Center), funded by the Research Foundation - Flanders (FWO) and the Flemish Government.

