## Supplementary material for "Open modification searching of SARS-CoV-2–human protein interaction data reveals novel viral modification sites": Supplementary Figures.pdf

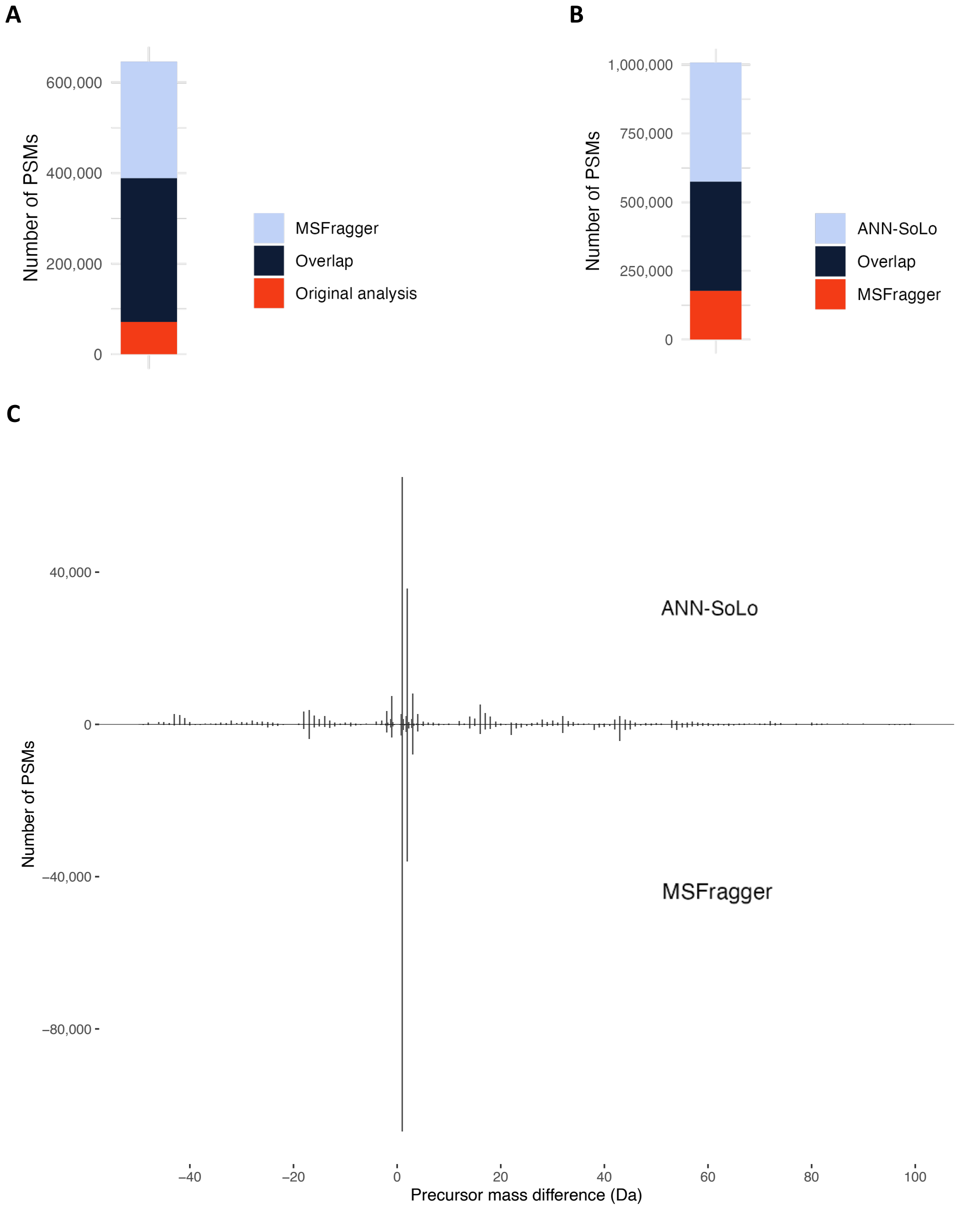


**Supplementary Figure 1. (A)** Comparison in identification performance between the originally reported results using standard searching and open modification searching with MSFragger. Open modification searching was able to identify more spectra, corresponding to the identification of a large number of modified PSMs. **(B)** Comparison of the identification results from ANN-SoLo with the identification results of MSFragger. **(C)** Mirror plot illustrating the precursor mass differences observed by open modification searching with ANN-SoLo (top) and MSFragger (bottom).

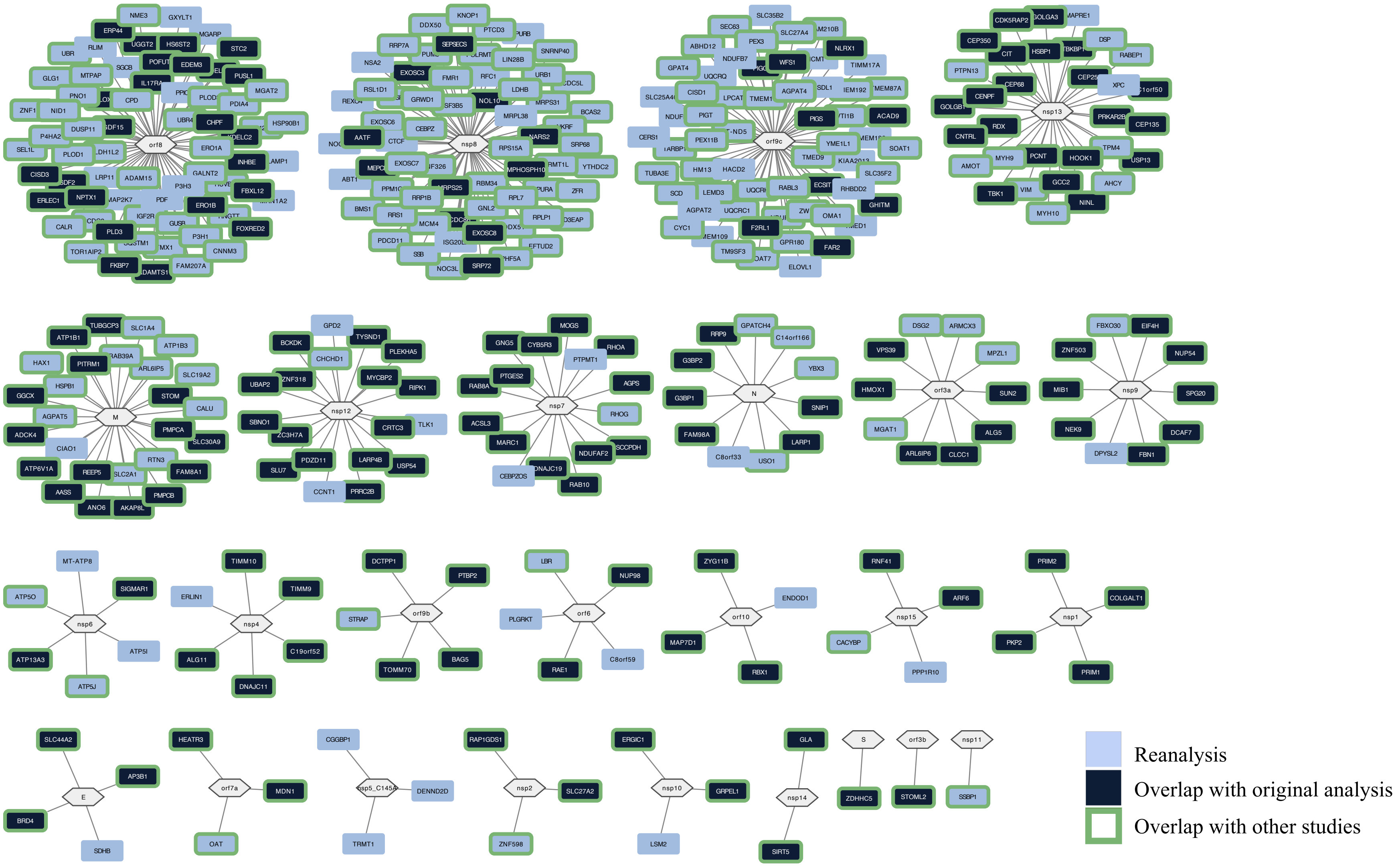


**Supplementary Figure 2.** Interaction network of the 375 identified PPIs, including 164 interactions that overlap in the original study, and 124 interactions that were found in independent SARS-CoV-2 interactome studies.

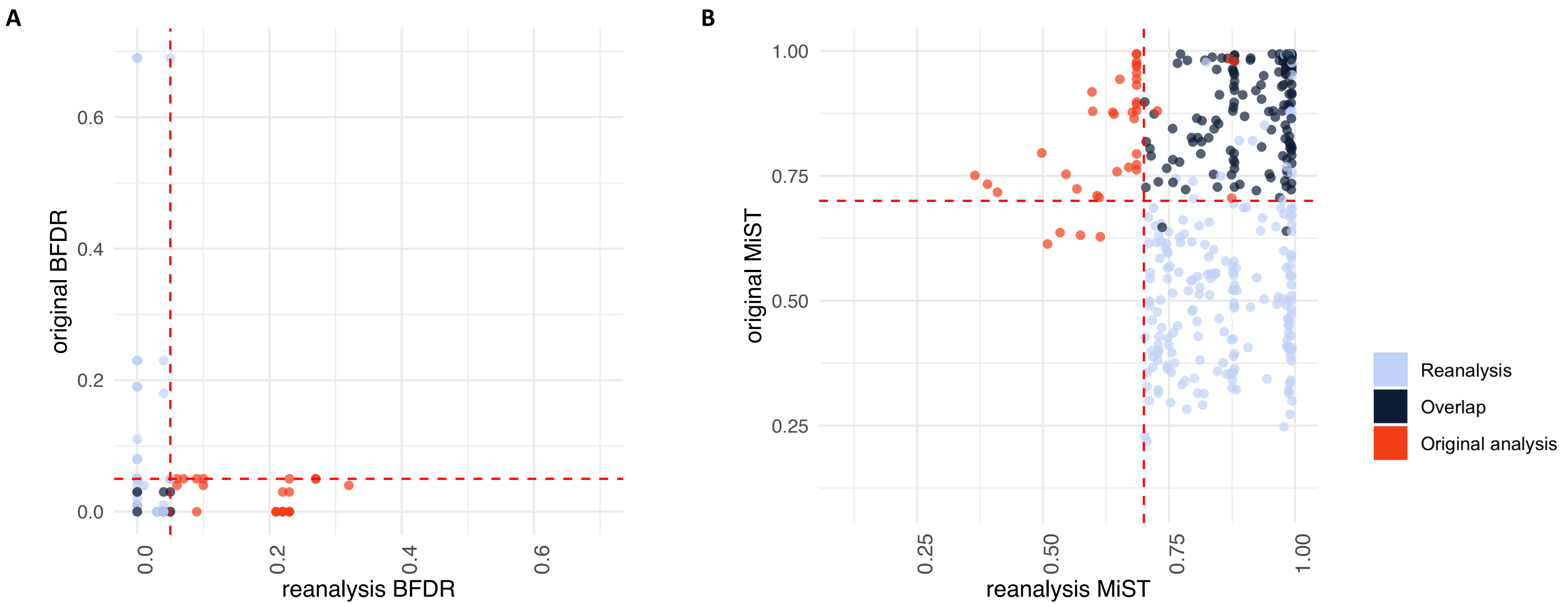


**Supplementary Figure 3.** 363 of the 375 PPIs identified in the reanalysis are present in the unfiltered results of the original analysis. 203 of the 332 identified PPIs in the original analysis are present in the unfiltered results of the reanalysis. Here we show **(A)** the SAINTexpress BFDR and **(B)** the MiST scores of the original analysis plotted against those of the reanalysis for 402 identified PPIs present in both unfiltered results. The red dotted lines indicate the filtering thresholds used for the first step in the original two-step filtering strategy, which is the same as the thresholds used during the reanalysis (SAINTexpress BFDR $\leq 0.05$, MiST score $\geq 0.7$).

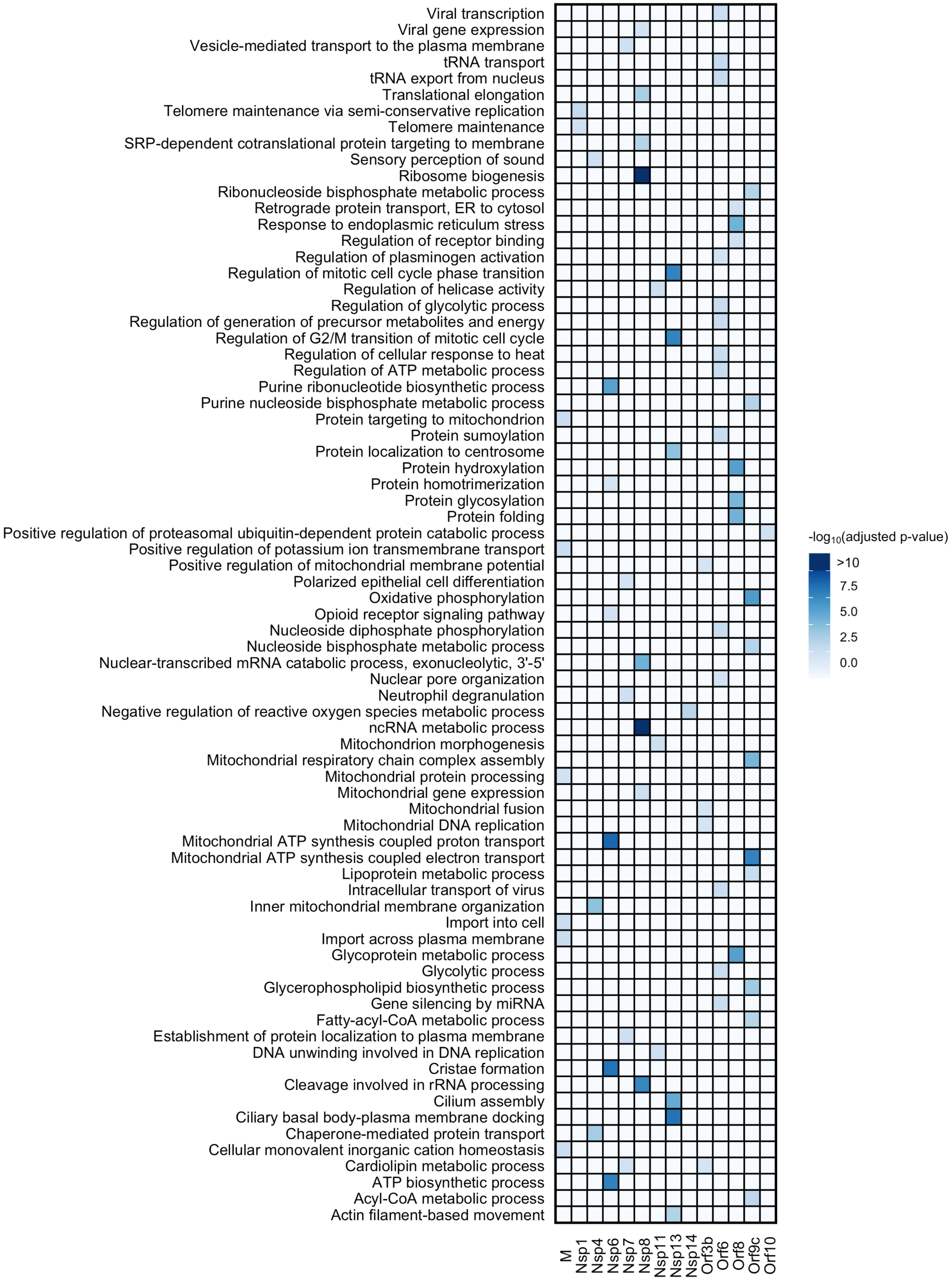


**Supplementary Figure 4.** GO enrichment analysis for each of the SARS-CoV-2 proteins.

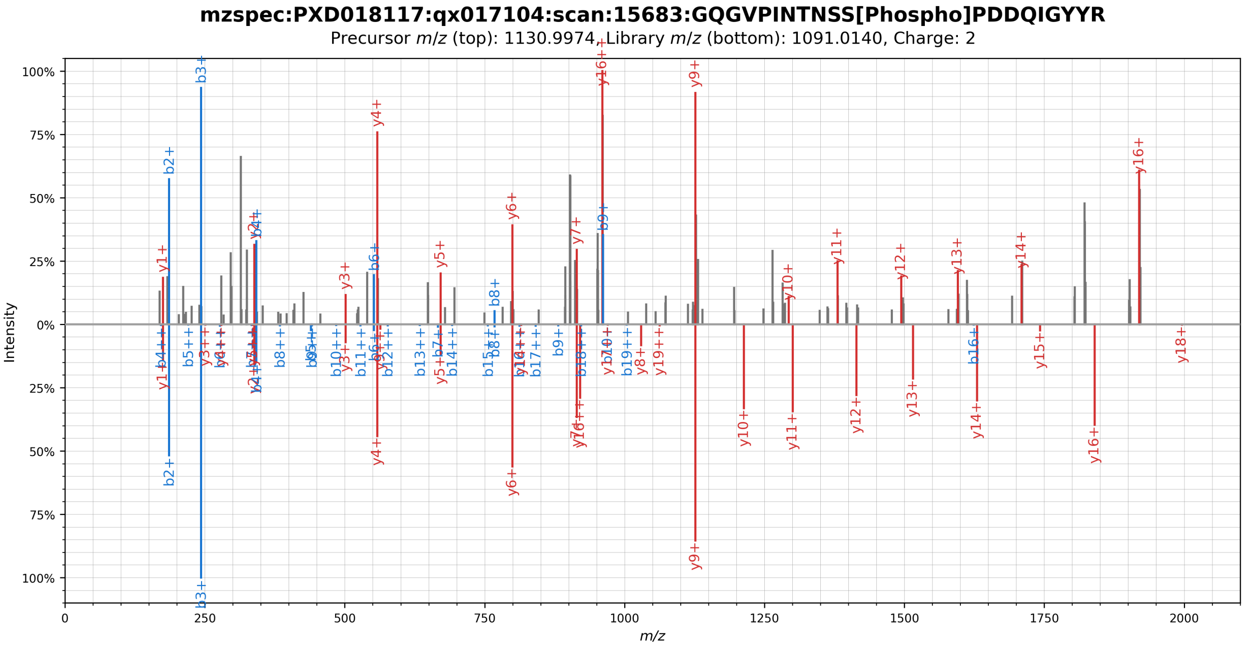


**Supplementary Figure 5.** Mirror plot showing the spectral library match (bottom) of a phosphorylated peptide originating from the SARS-CoV-2 N protein (top). The phosphorylation was localized to residue S79 by visual inspection.

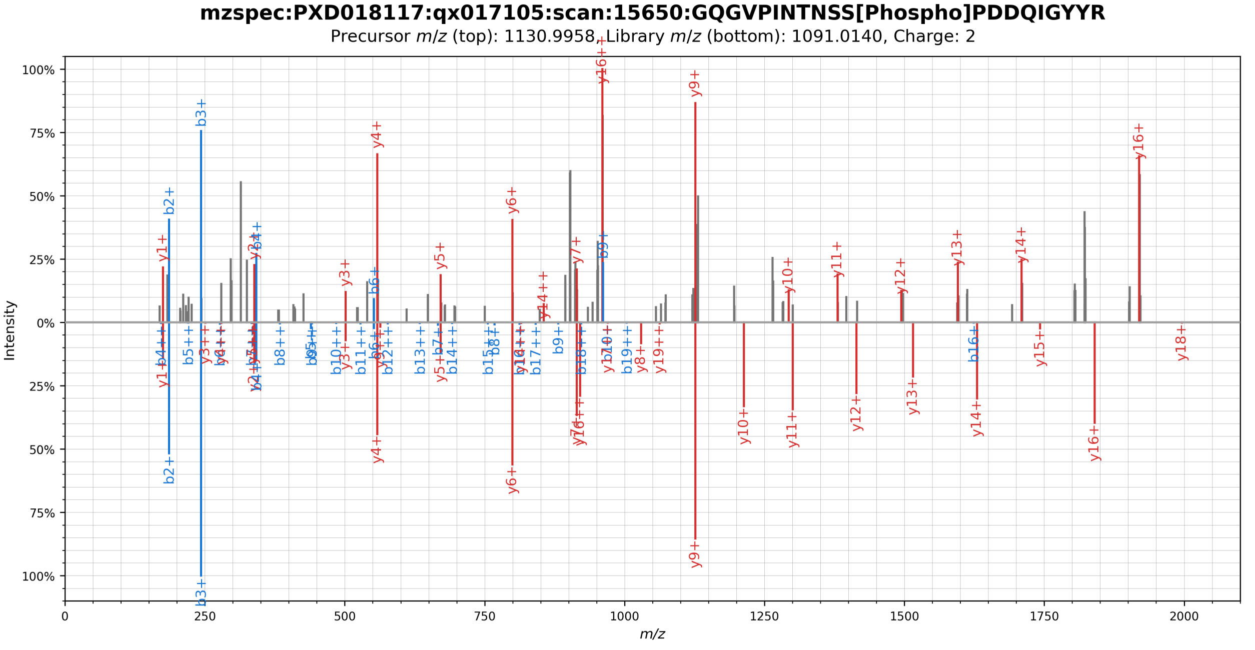


**Supplementary Figure 6.** Mirror plot showing the spectral library match (bottom) of a phosphorylated peptide originating from the SARS-CoV-2 N protein (top). The phosphorylation was localized to residue S79 by visual inspection.

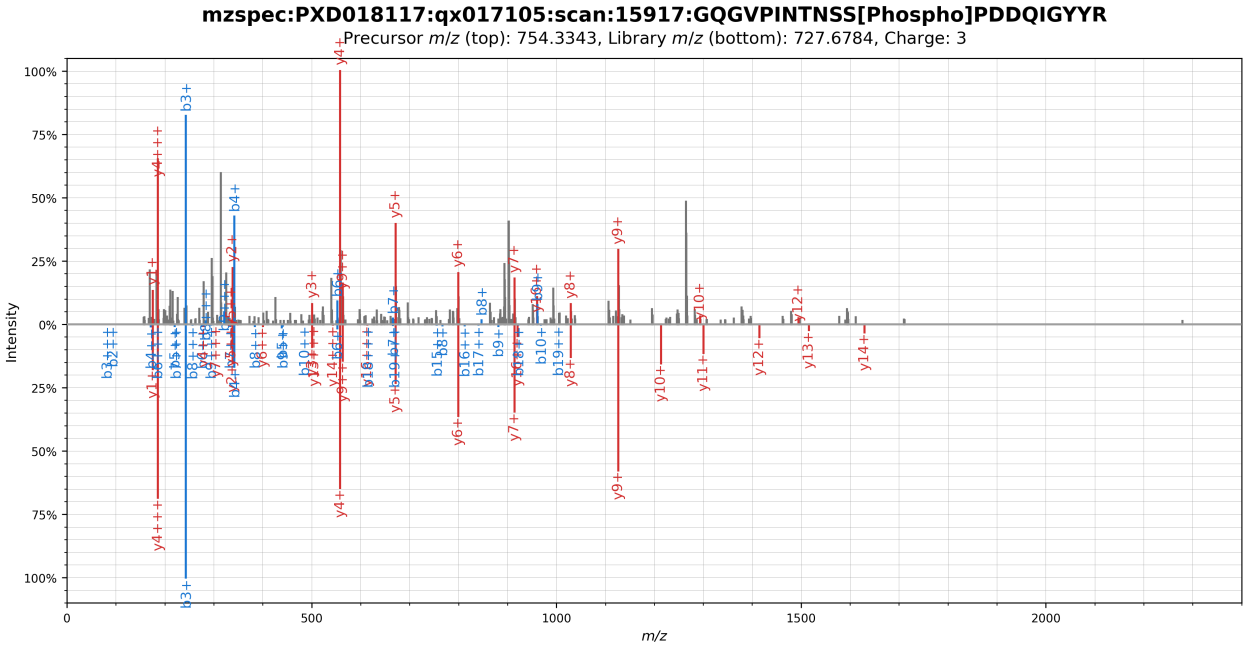


**Supplementary Figure 7.** Mirror plot showing the spectral library match (bottom) of a phosphorylated peptide originating from the SARS-CoV-2 N protein (top). The phosphorylation was localized to residue S79 by visual inspection.

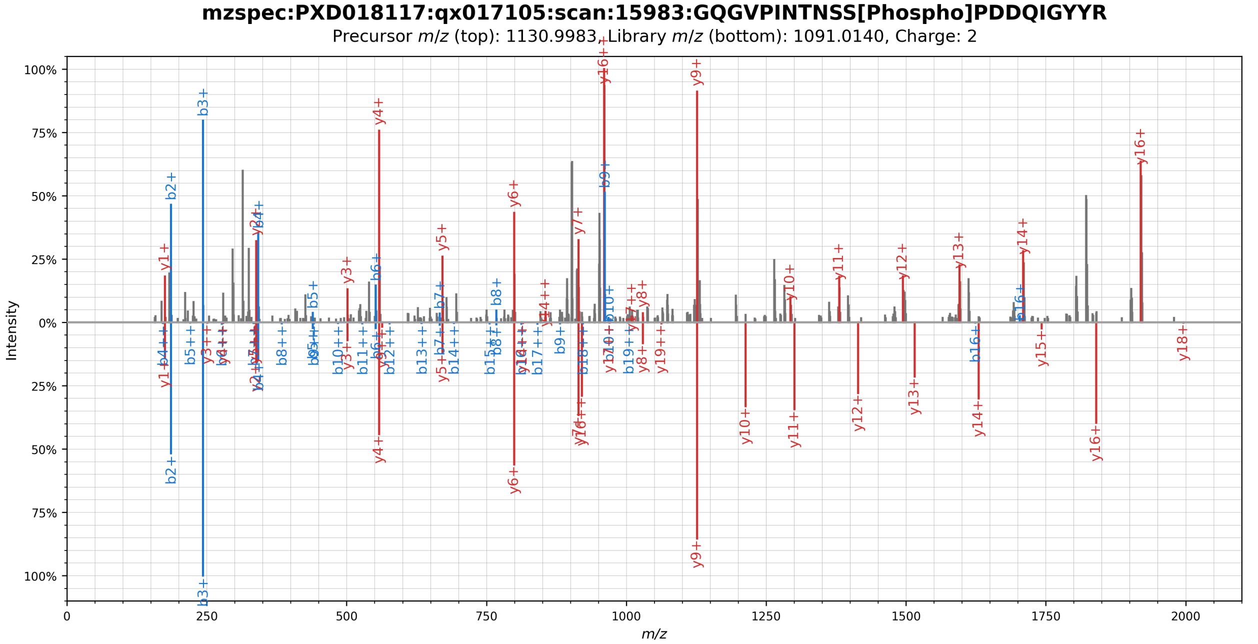


**Supplementary Figure 8.** Mirror plot showing the spectral library match (bottom) of a phosphorylated peptide originating from the SARS-CoV-2 N protein (top). The phosphorylation was localized to residue S79 by visual inspection.

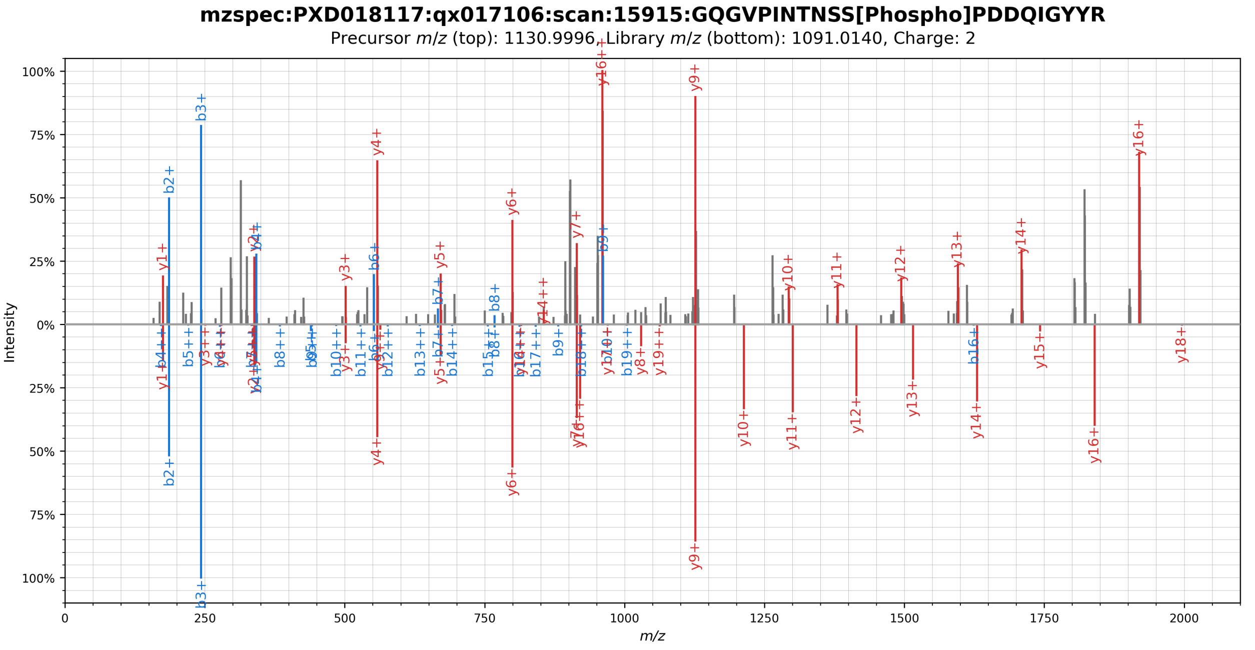


**Supplementary Figure 9.** Mirror plot showing the spectral library match (bottom) of a phosphorylated peptide originating from the SARS-CoV-2 N protein (top). The phosphorylation was localized to residue S79 by visual inspection.

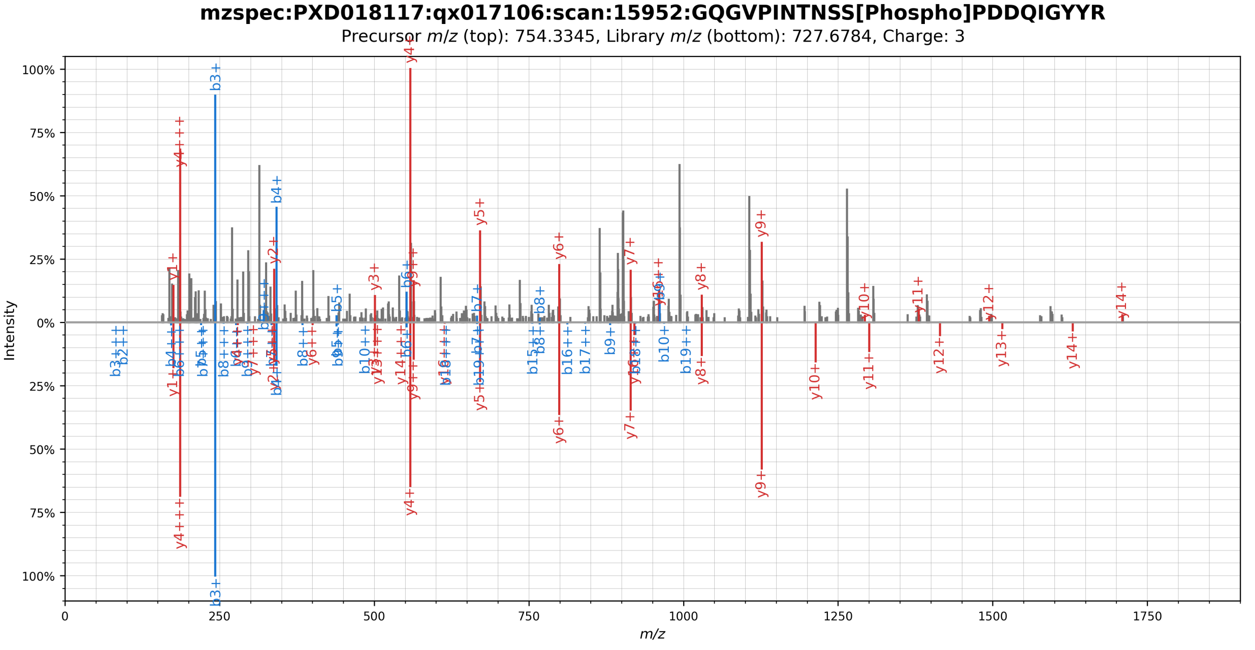


**Supplementary Figure 10.** Mirror plot showing the spectral library match (bottom) of a phosphorylated peptide originating from the SARS-CoV-2 N protein (top). The phosphorylation was localized to residue S79 by visual inspection.

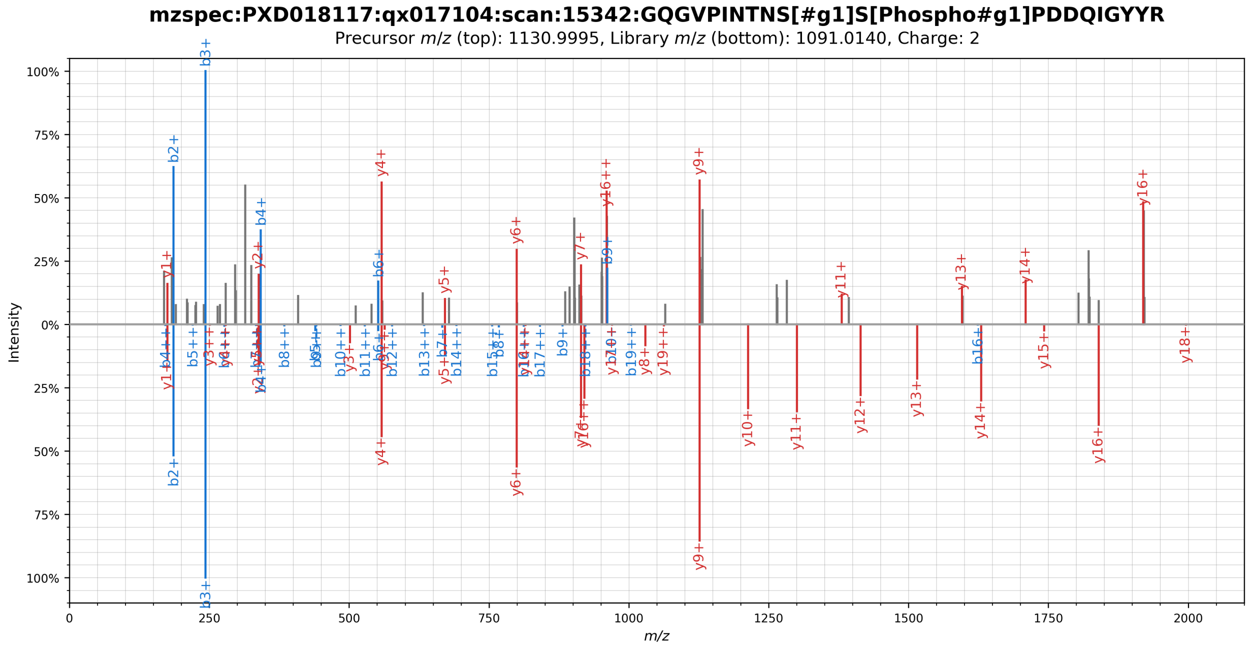


**Supplementary Figure 11.** Mirror plot showing the spectral library match (bottom) of a phosphorylated peptide originating from the SARS-CoV-2 N protein (top). The phosphorylation was localized to residue S78/S79 by visual inspection.

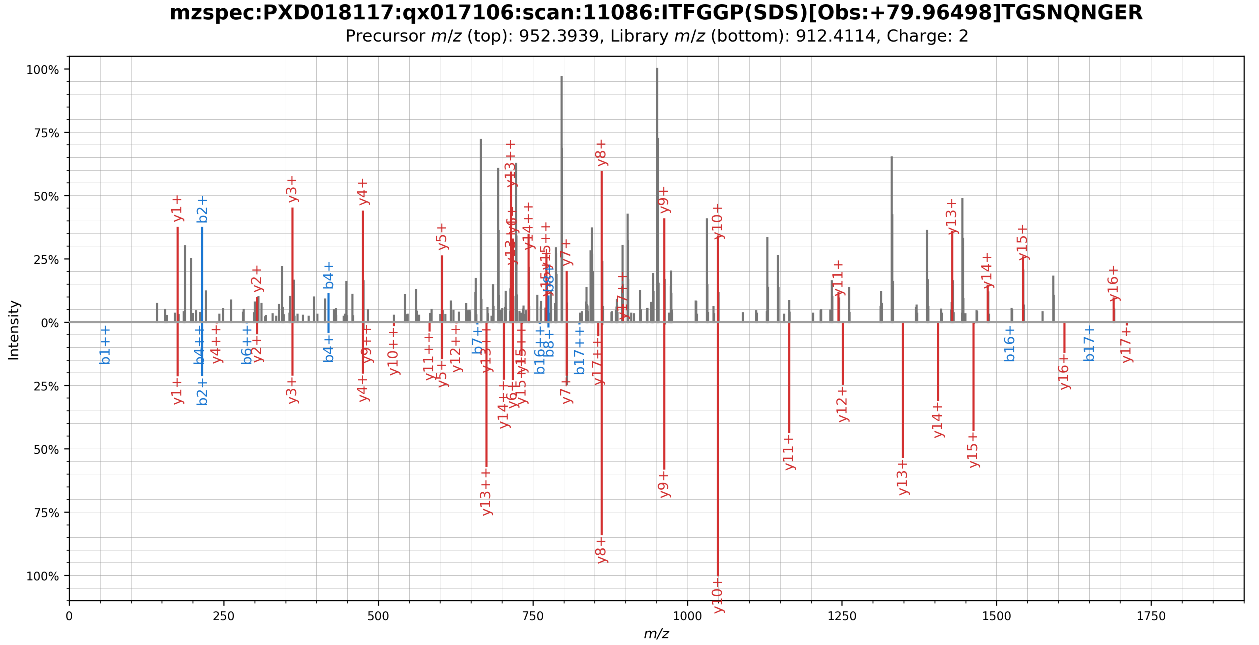


**Supplementary Figure 12.** Mirror plot showing the spectral library match (bottom) of a phosphorylated peptide originating from the SARS-CoV-2 N protein (top). The phosphorylation was localized to residue S21/S23 by visual inspection.

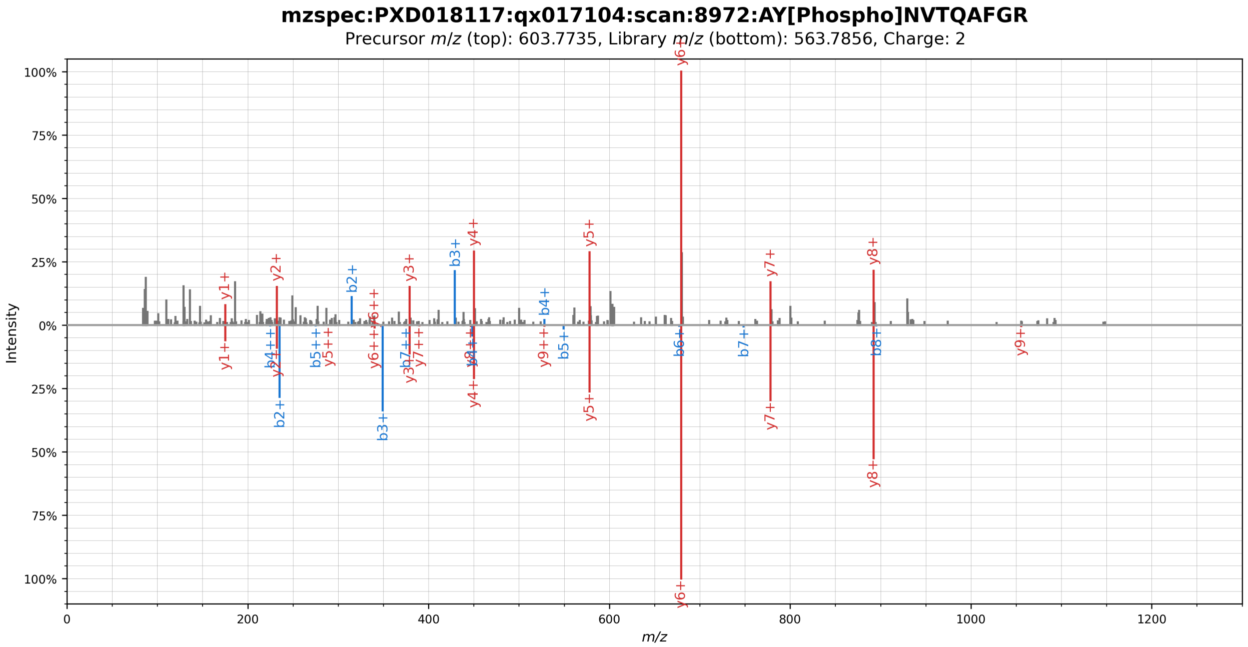


**Supplementary Figure 13.** Mirror plot showing the spectral library match (bottom) of a phosphorylated peptide originating from the SARS-CoV-2 N protein (top). The phosphorylation was localized to residue Y268 by visual inspection.

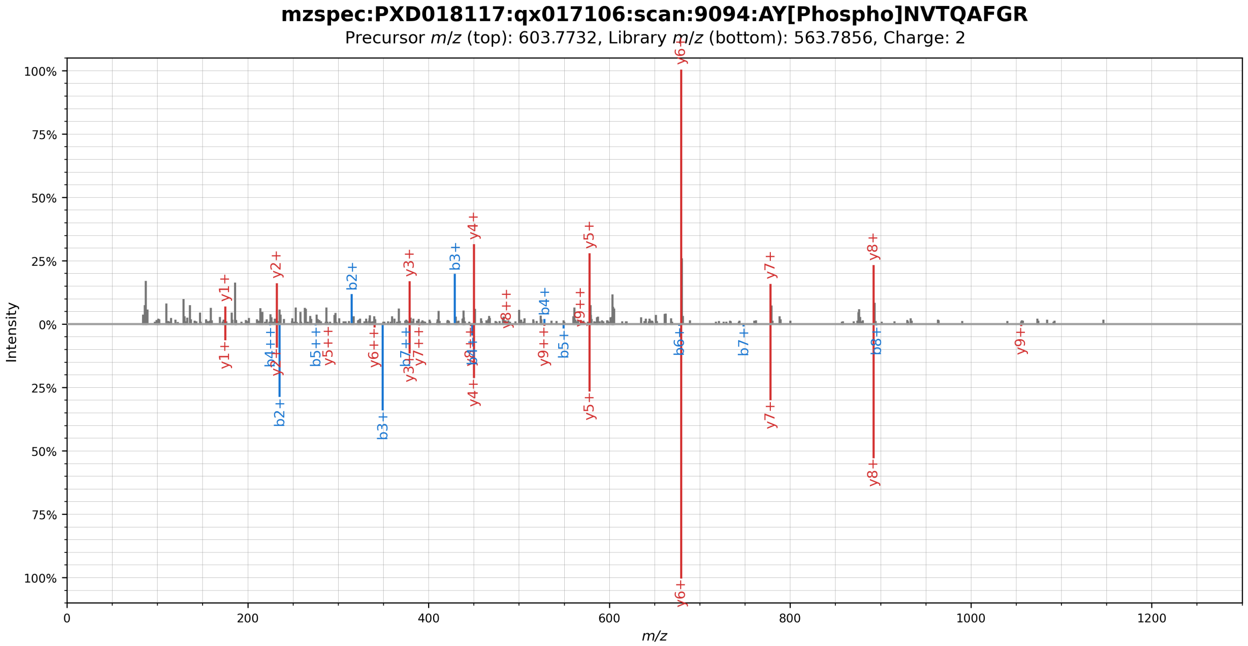


**Supplementary Figure 14.** Mirror plot showing the spectral library match (bottom) of a phosphorylated peptide originating from the SARS-CoV-2 N protein (top). The phosphorylation was localized to residue Y268 by visual inspection.

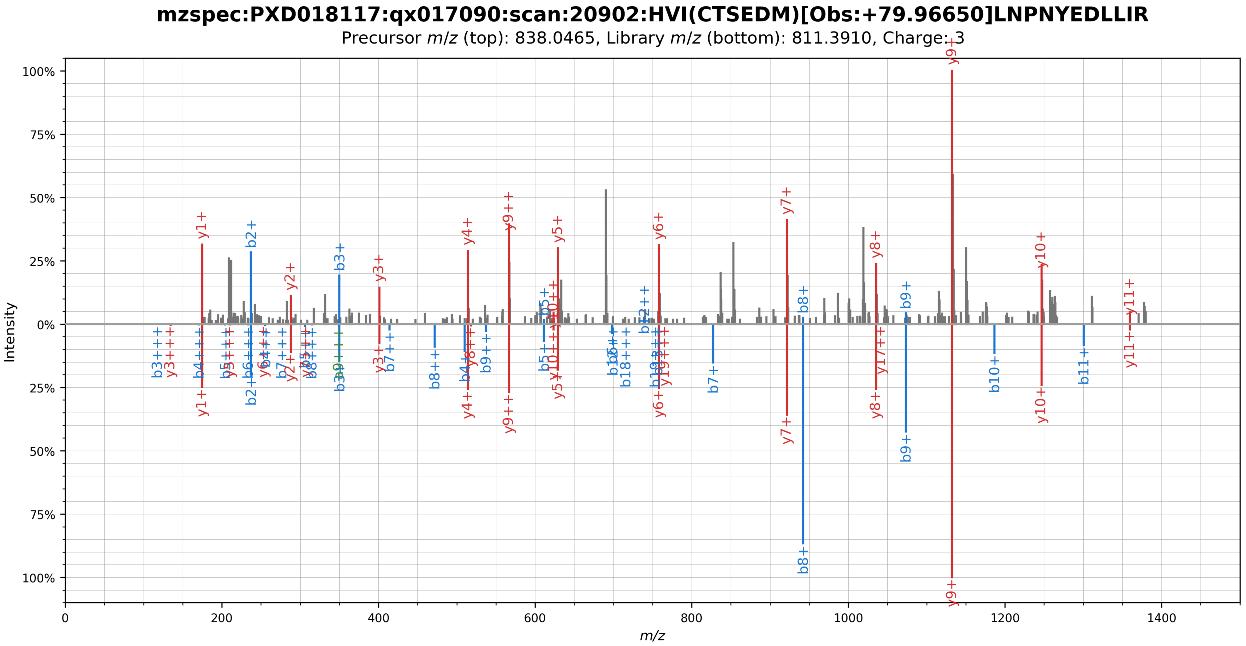


**Supplementary Figure 15.** Mirror plot showing the spectral library match (bottom) of a phosphorylated peptide originating from the SARS-CoV-2 Nsp5 protein (top). The phosphorylation was localized to residue T46/S47 by visual inspection.

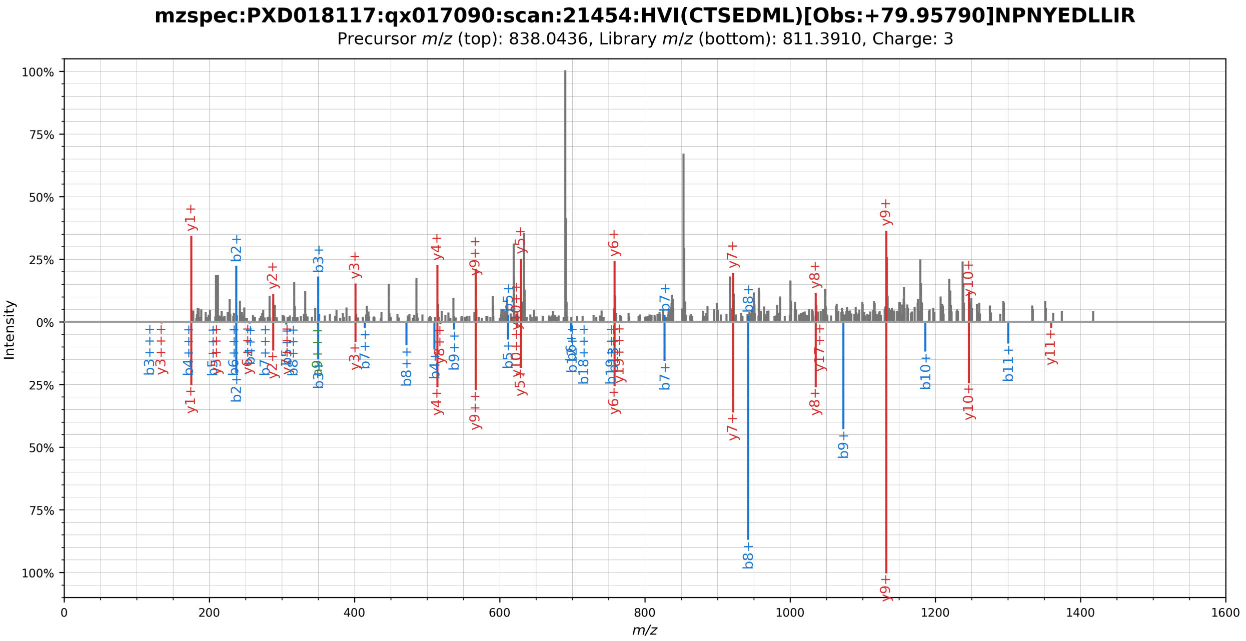


**Supplementary Figure 16.** Mirror plot showing the spectral library match (bottom) of a phosphorylated peptide originating from the SARS-CoV-2 Nsp5 protein (top). The phosphorylation was localized to residue T46/S47 by visual inspection.

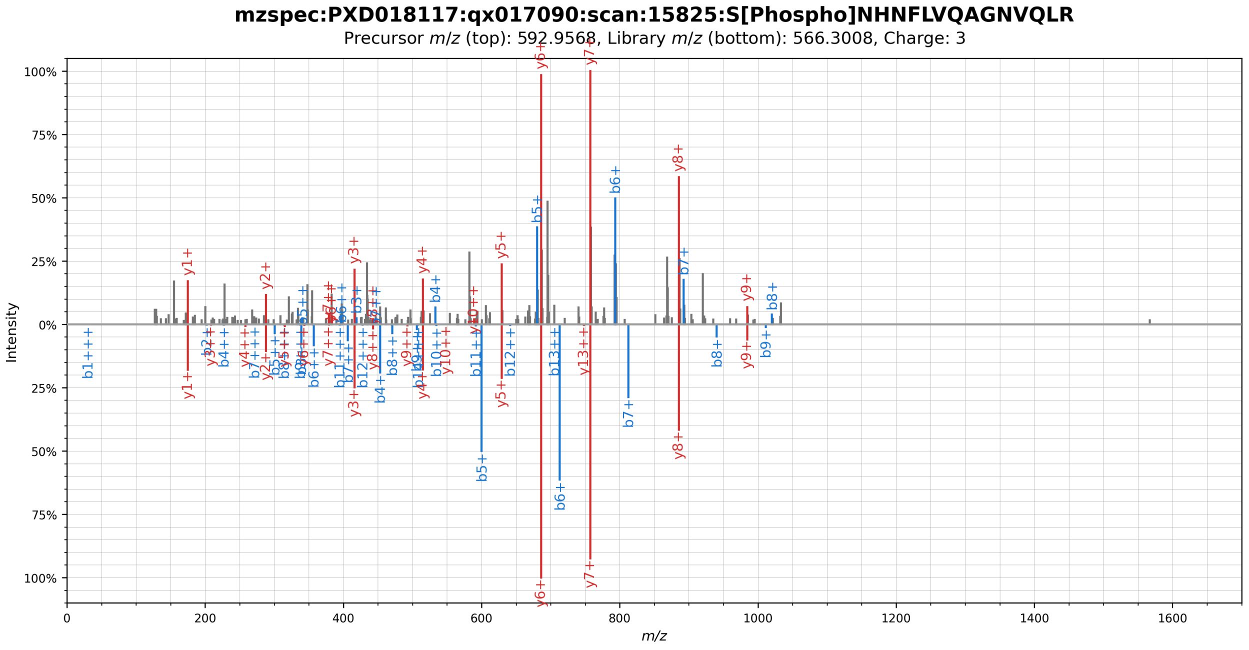


**Supplementary Figure 17.** Mirror plot showing the spectral library match (bottom) of a phosphorylated peptide originating from the SARS-CoV-2 Nsp5 protein (top). The phosphorylation was localized to residue S63 by visual inspection.

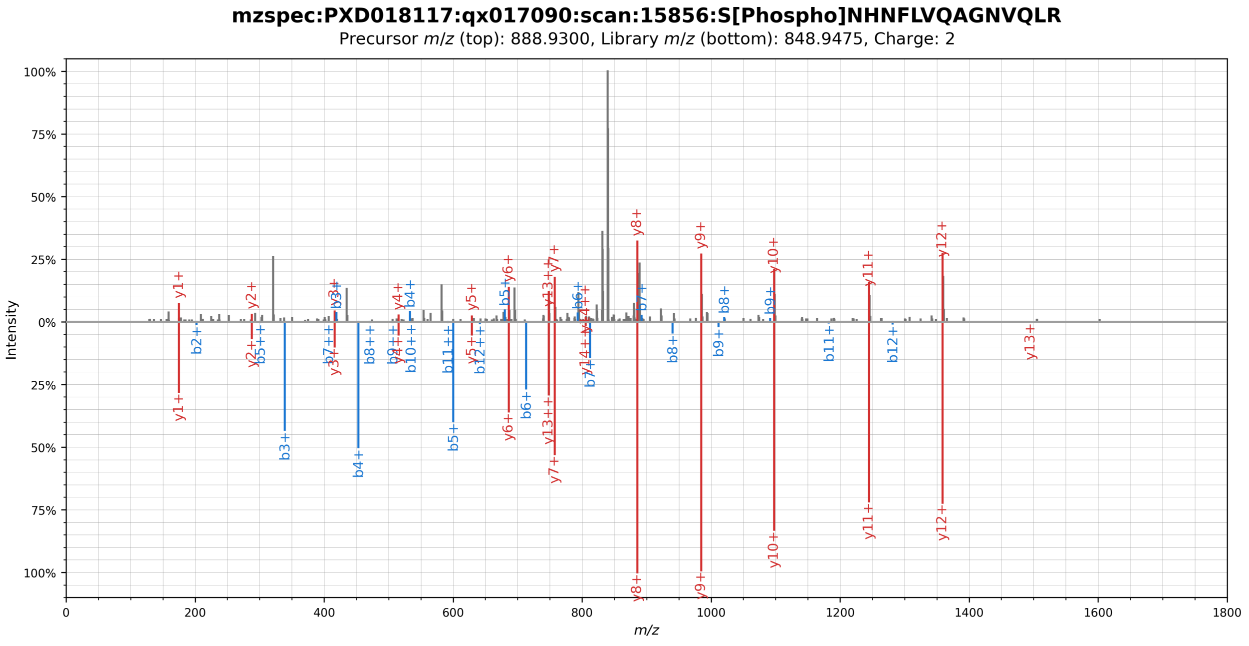


**Supplementary Figure 18.** Mirror plot showing the spectral library match (bottom) of a phosphorylated peptide originating from the SARS-CoV-2 Nsp5 protein (top). The phosphorylation was localized to residue S63 by visual inspection.

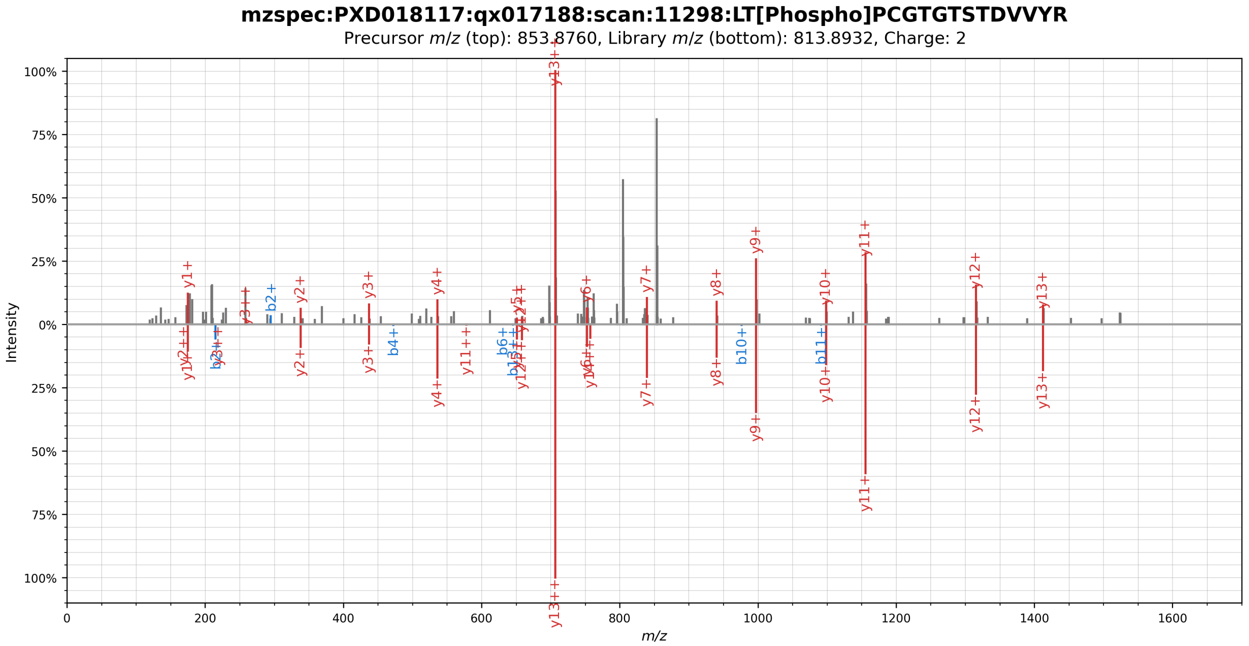


**Supplementary Figure 19.** Mirror plot showing the spectral library match (bottom) of a phosphorylated peptide originating from the SARS-CoV-2 Nsp12 protein (top). The phosphorylation was localized to residue T21 by visual inspection.

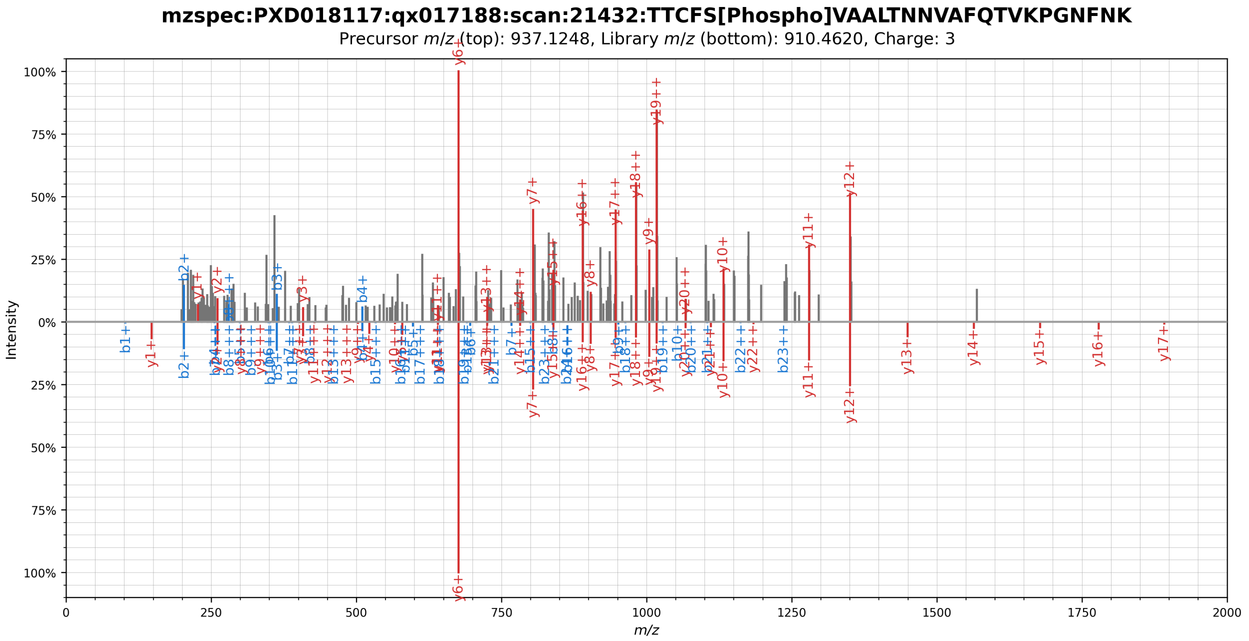


**Supplementary Figure 20.** Mirror plot showing the spectral library match (bottom) of a phosphorylated peptide originating from the SARS-CoV-2 Nsp12 protein (top). The phosphorylation was localized to residue S398 by visual inspection.

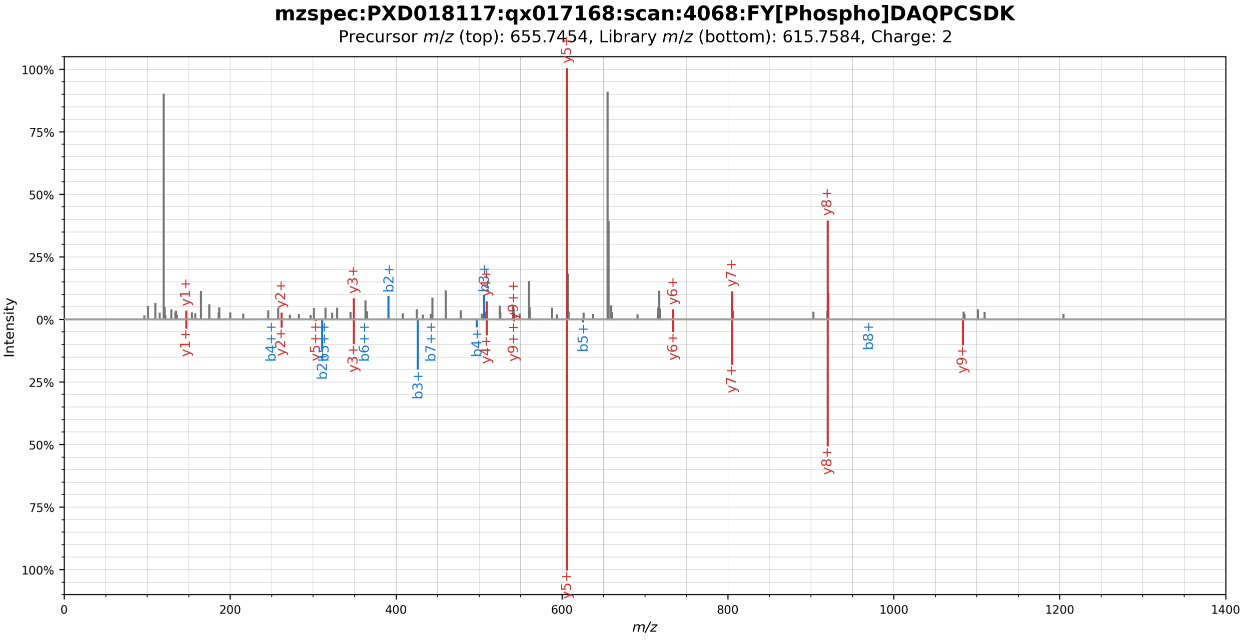


**Supplementary Figure 21.** Mirror plot showing the spectral library match (bottom) of a phosphorylated peptide originating from the SARS-CoV-2 Nsp14 protein (top). The phosphorylation was localized to residue Y351 by visual inspection.

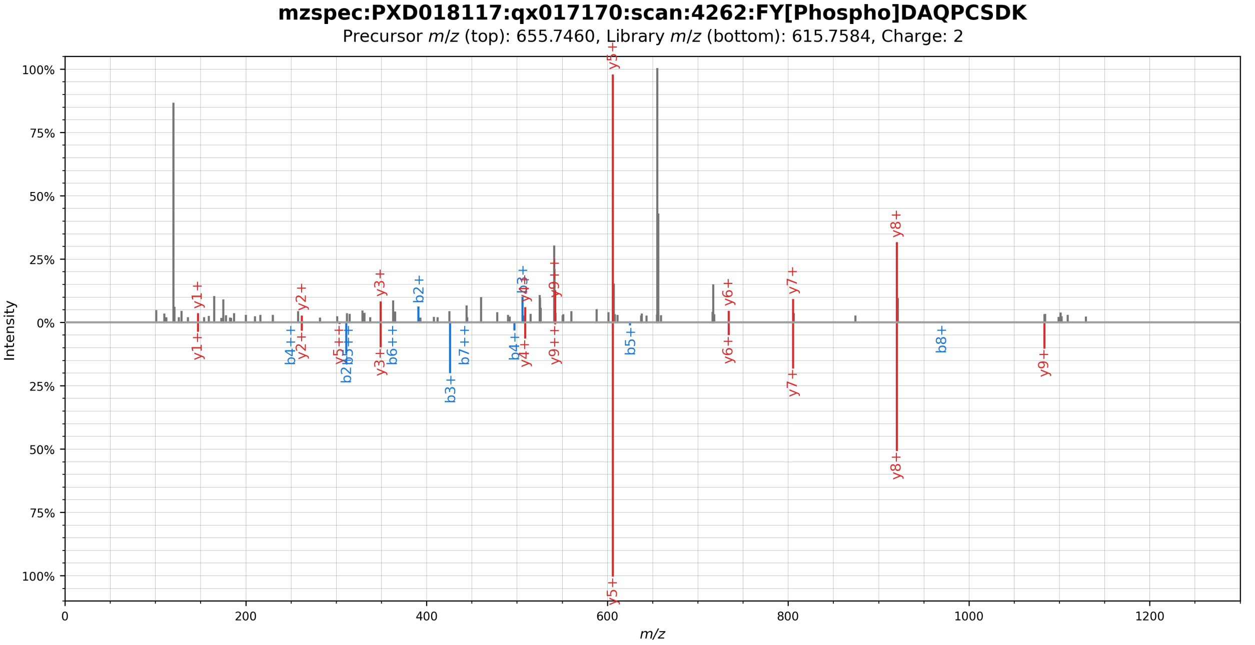


**Supplementary Figure 22.** Mirror plot showing the spectral library match (bottom) of a phosphorylated peptide originating from the SARS-CoV-2 Nsp14 protein (top). The phosphorylation was localized to residue Y351 by visual inspection.

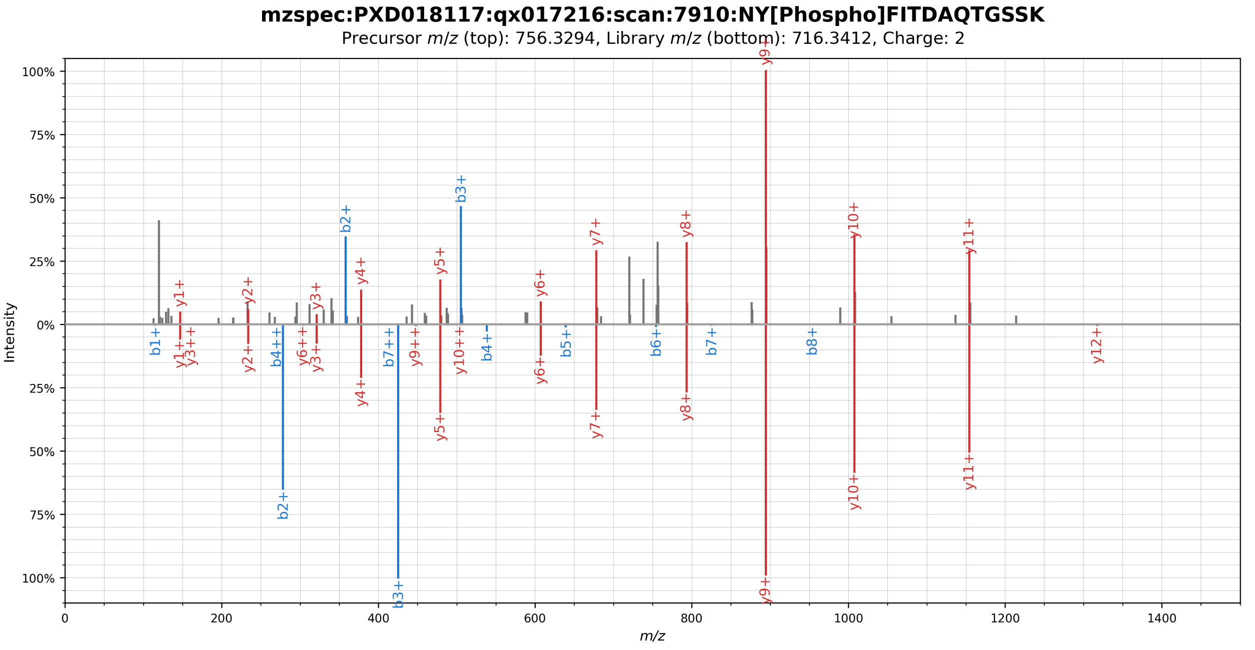


**Supplementary Figure 23.** Mirror plot showing the spectral library match (bottom) of a phosphorylated peptide originating from the SARS-CoV-2 Nsp15 protein (top). The phosphorylation was localized to residue Y279 by visual inspection.

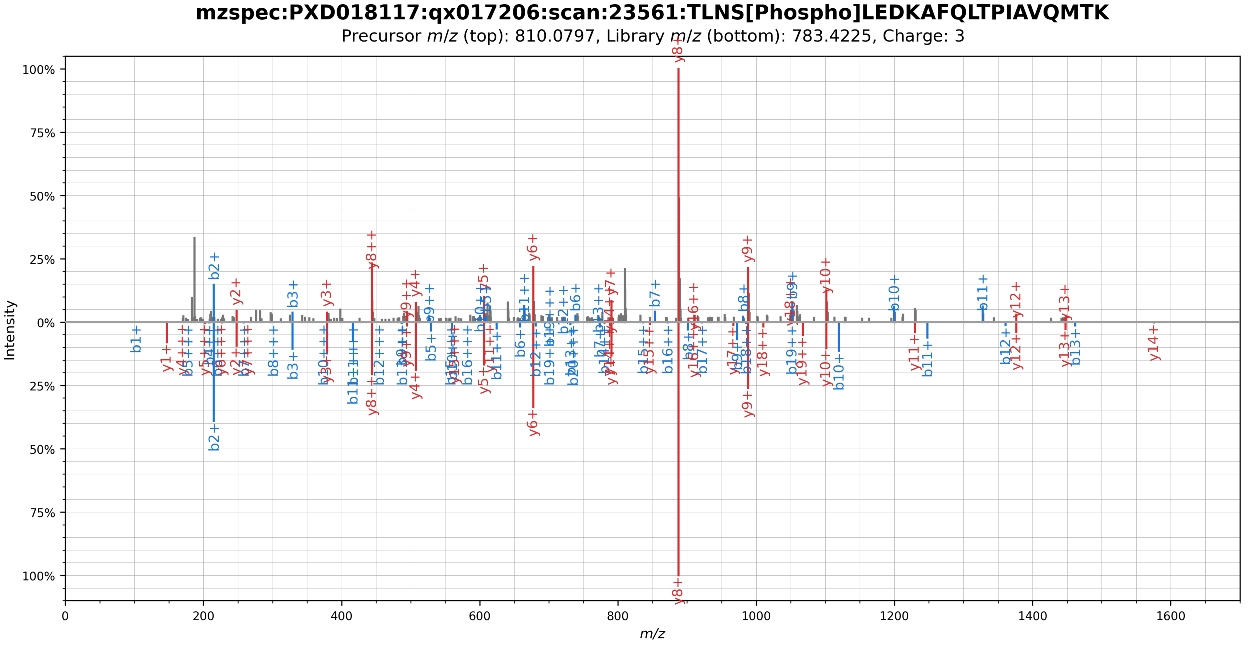


**Supplementary Figure 24.** Mirror plot showing the spectral library match (bottom) of a phosphorylated peptide originating from the SARS-CoV-2 Orf9b protein (top). The phosphorylation was localized to residue S63 by visual inspection.

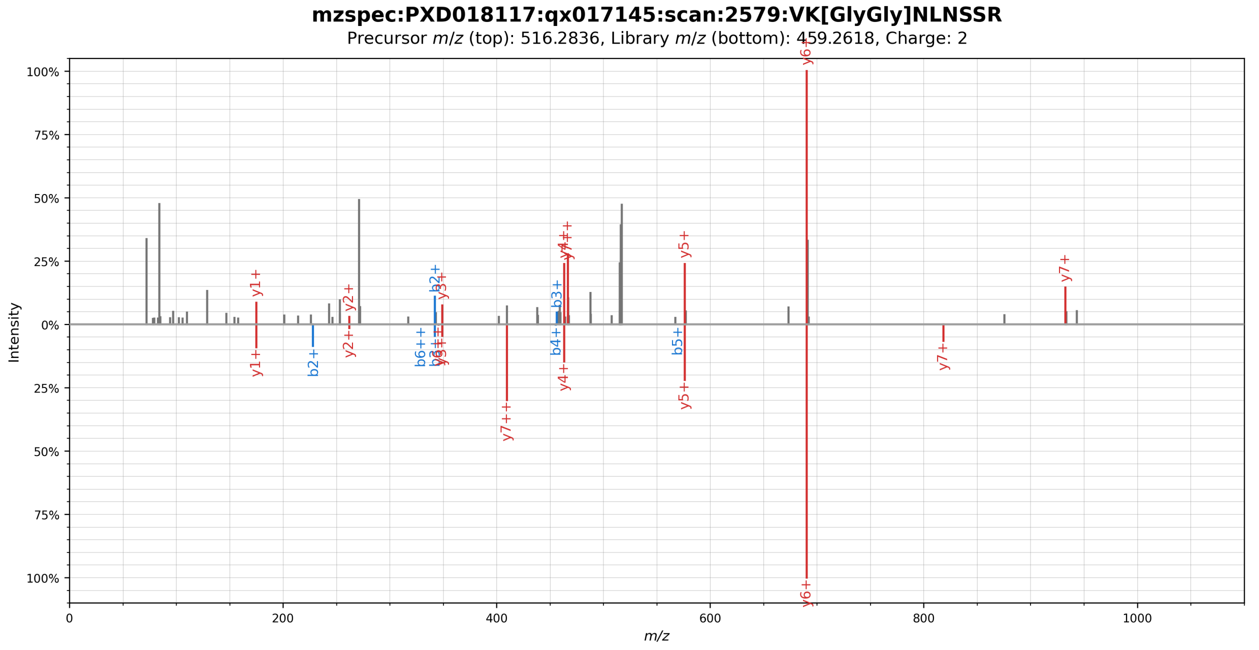


**Supplementary Figure 25.** Mirror plot showing the spectral library match (bottom) of a ubiquitylated peptide originating from the SARS-CoV-2 E protein (top). The phosphorylation was localized to residue K63 by visual inspection.

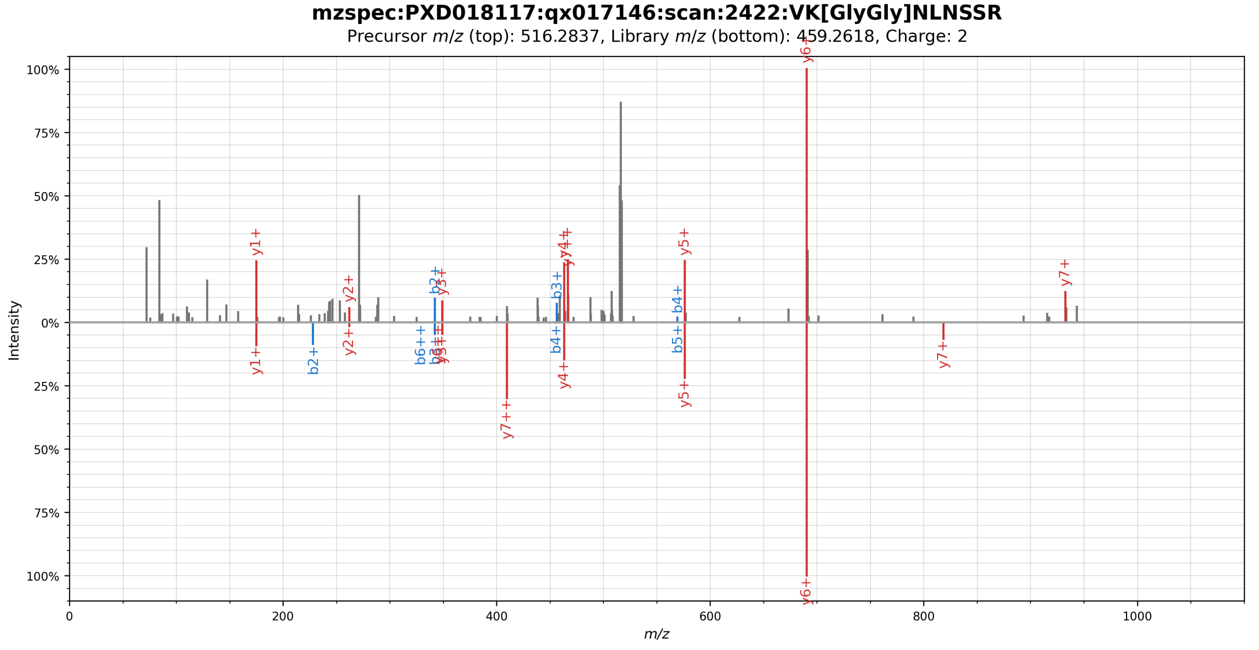


**Supplementary Figure 26.** Mirror plot showing the spectral library match (bottom) of a ubiquitylated peptide originating from the SARS-CoV-2 E protein (top). The phosphorylation was localized to residue K63 by visual inspection.

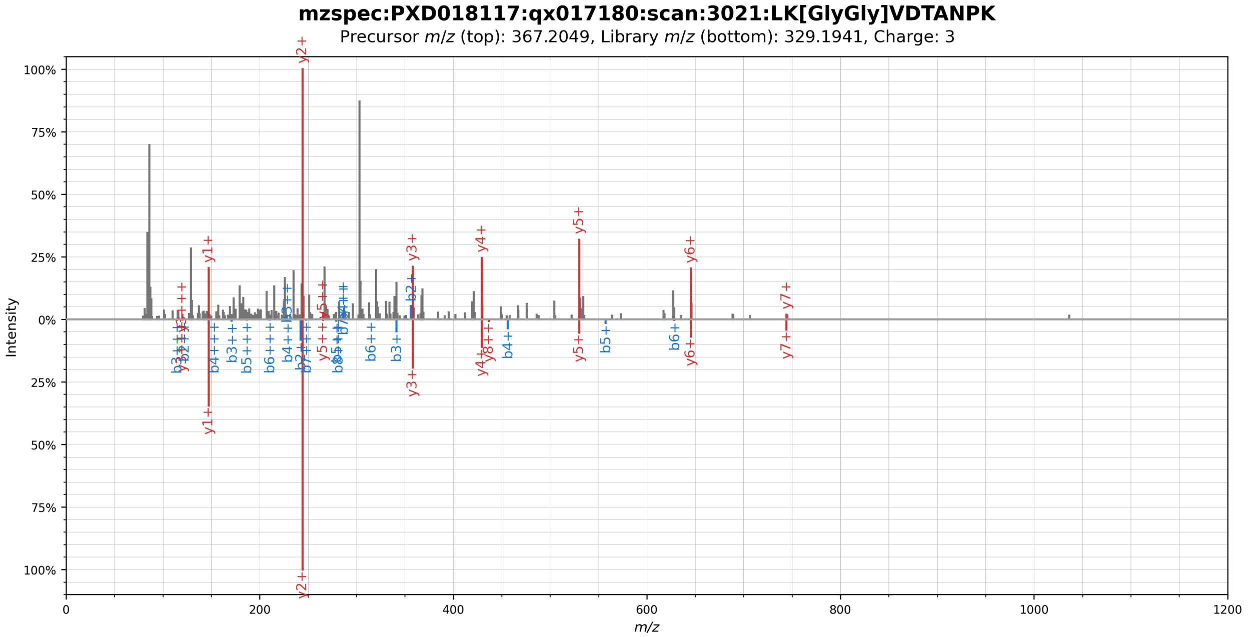


**Supplementary Figure 27.** Mirror plot showing the spectral library match (bottom) of a ubiquitylated peptide originating from the SARS-CoV-2 Nsp5(C145A) protein (top). The phosphorylation was localized to residue K90 by visual inspection.

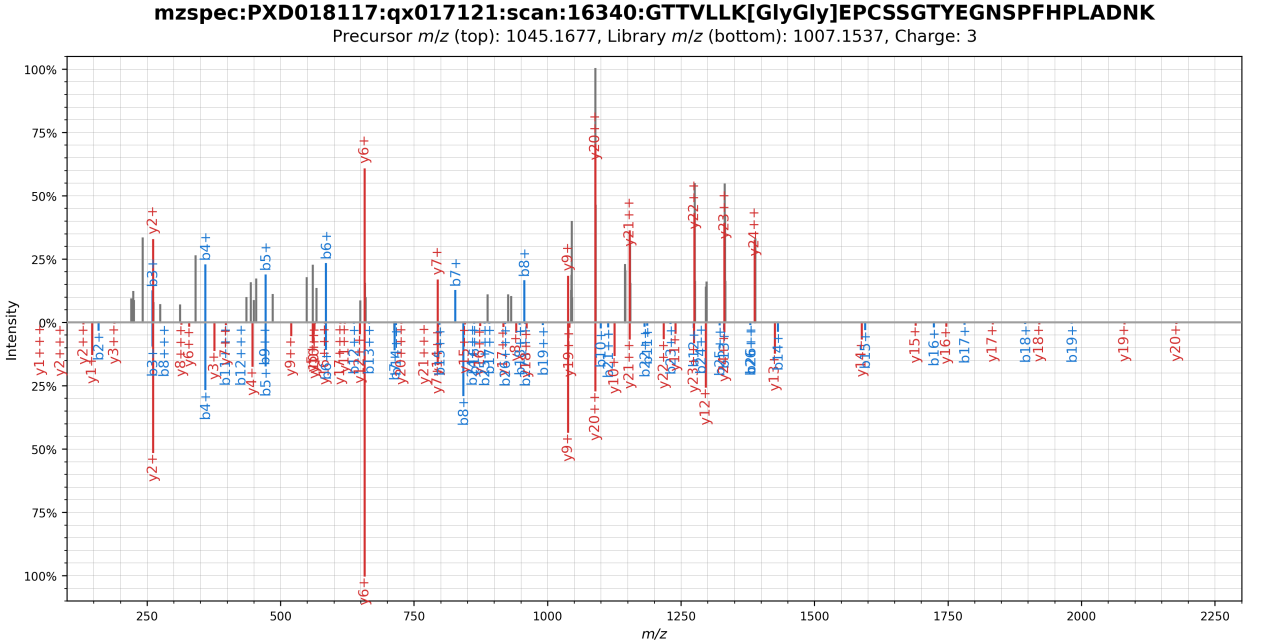


**Supplementary Figure 28.** Mirror plot showing the spectral library match (bottom) of a ubiquitylated peptide originating from the SARS-CoV-2 Orf7a protein (top). The phosphorylation was localized to residue K32 by visual inspection.

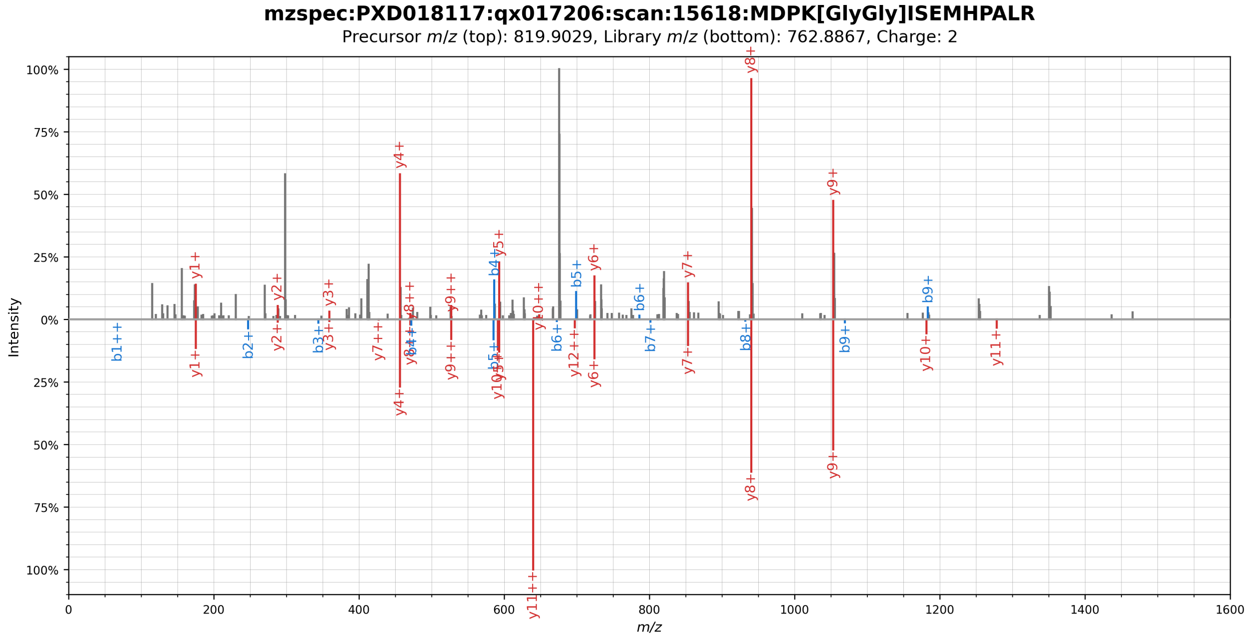


**Supplementary Figure 29.** Mirror plot showing the spectral library match (bottom) of a ubiquitylated peptide originating from the SARS-CoV-2 Orf9b protein (top). The phosphorylation was localized to residue K4 by visual inspection.

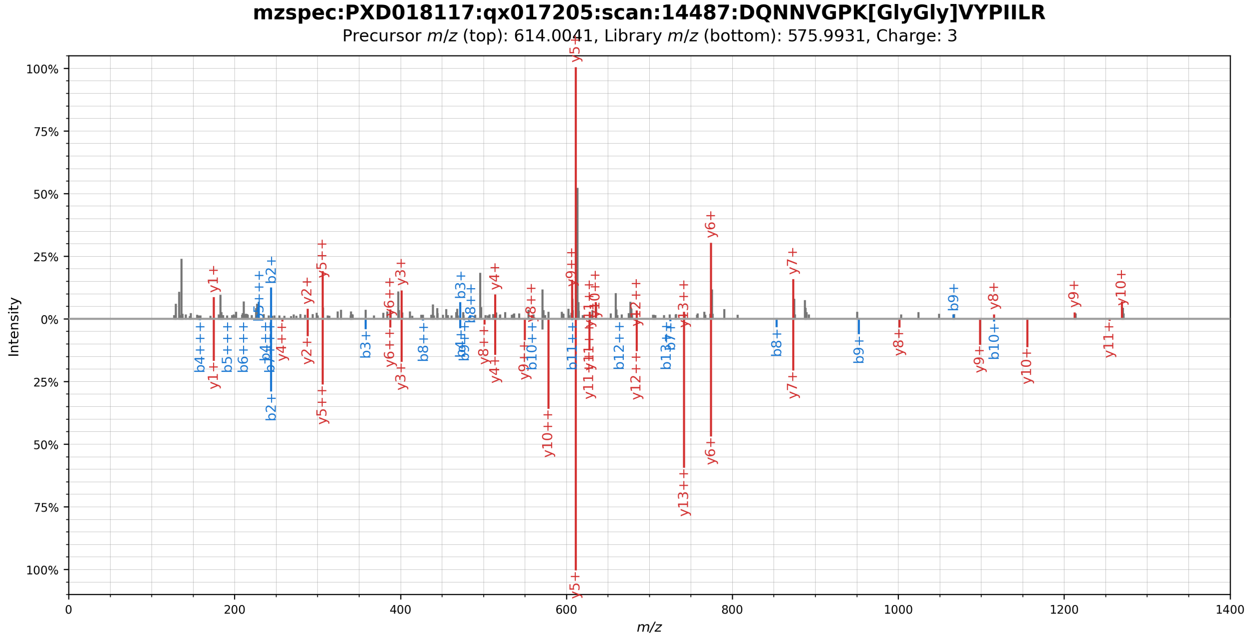


**Supplementary Figure 30.** Mirror plot showing the spectral library match (bottom) of a ubiquitylated peptide originating from the SARS-CoV-2 Orf9b protein (top). The phosphorylation was localized to residue K40 by visual inspection.

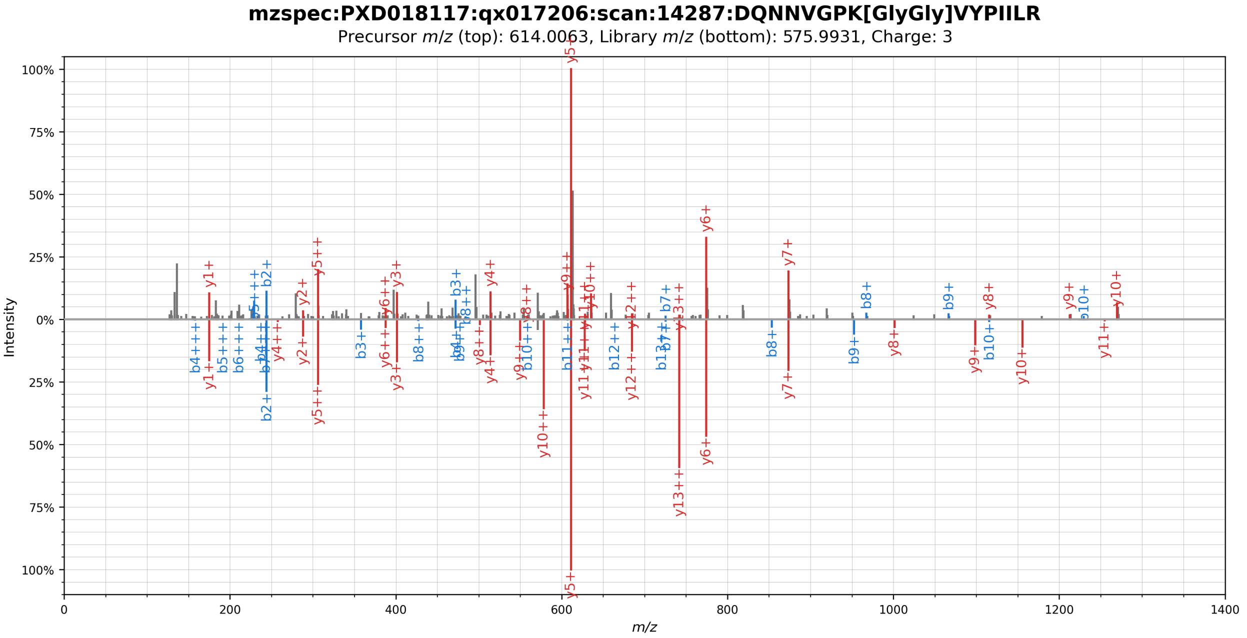


**Supplementary Figure 31.** Mirror plot showing the spectral library match (bottom) of a ubiquitylated peptide originating from the SARS-CoV-2 Orf9b protein (top). The phosphorylation was localized to residue K40 by visual inspection.

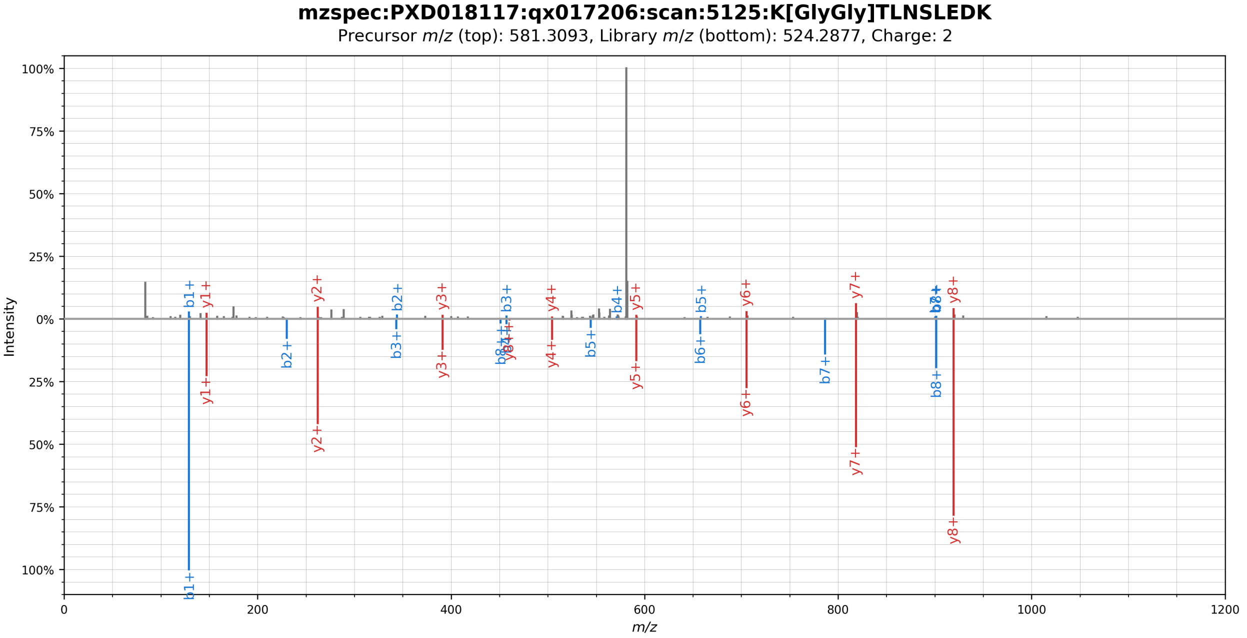


**Supplementary Figure 32.** Mirror plot showing the spectral library match (bottom) of a ubiquitylated peptide originating from the SARS-CoV-2 Orf9b protein (top). The phosphorylation was localized to residue K59 by visual inspection.

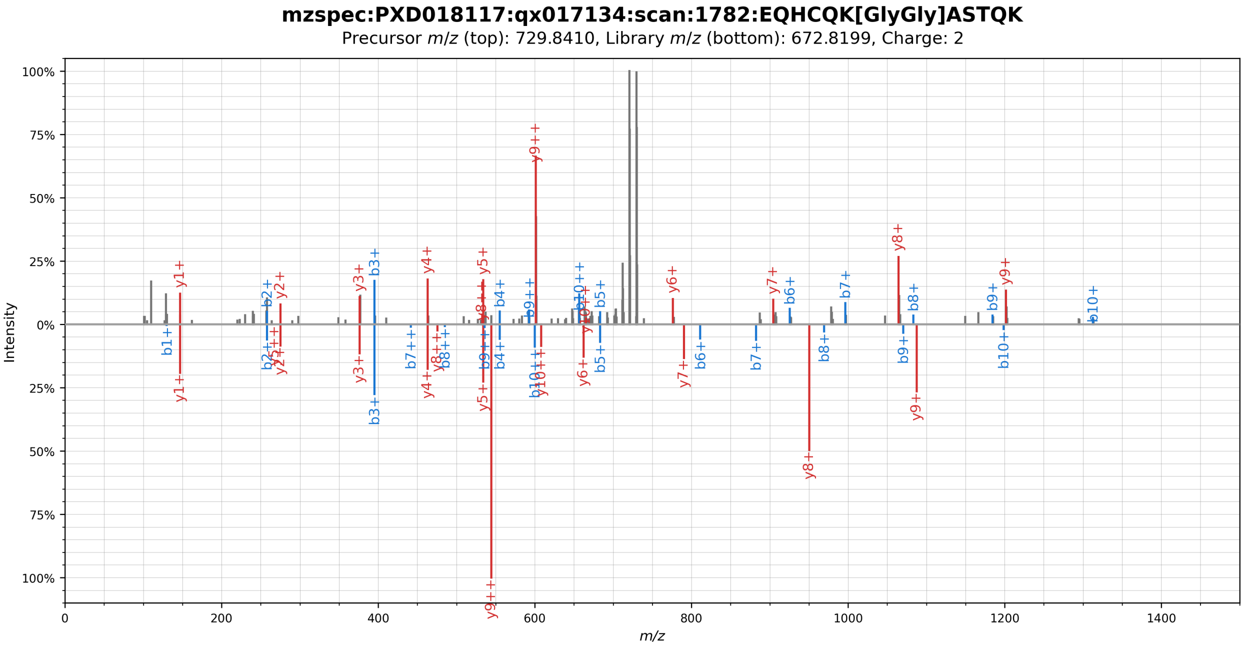


**Supplementary Figure 33.** Mirror plot showing the spectral library match (bottom) of a ubiquitylated peptide originating from the SARS-CoV-2 Orf9c protein (top). The phosphorylation was localized to residue K16 by visual inspection.

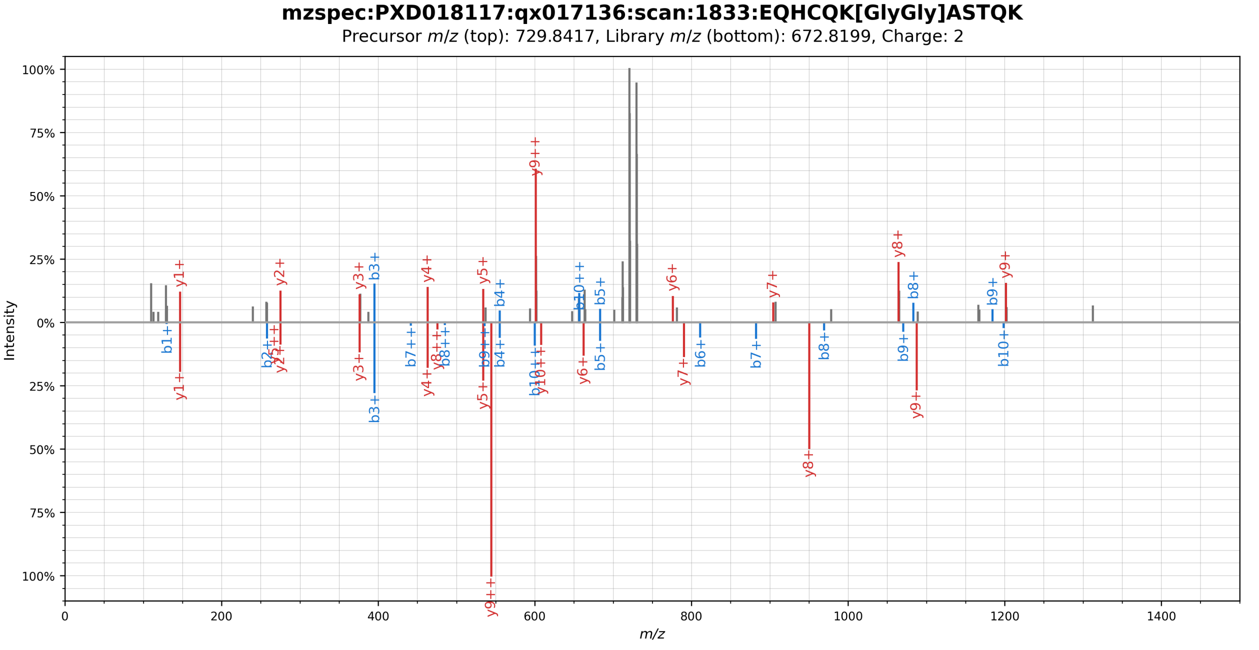


**Supplementary Figure 34.** Mirror plot showing the spectral library match (bottom) of a ubiquitylated peptide originating from the SARS-CoV-2 Orf9c protein (top). The phosphorylation was localized to residue K16 by visual inspection.

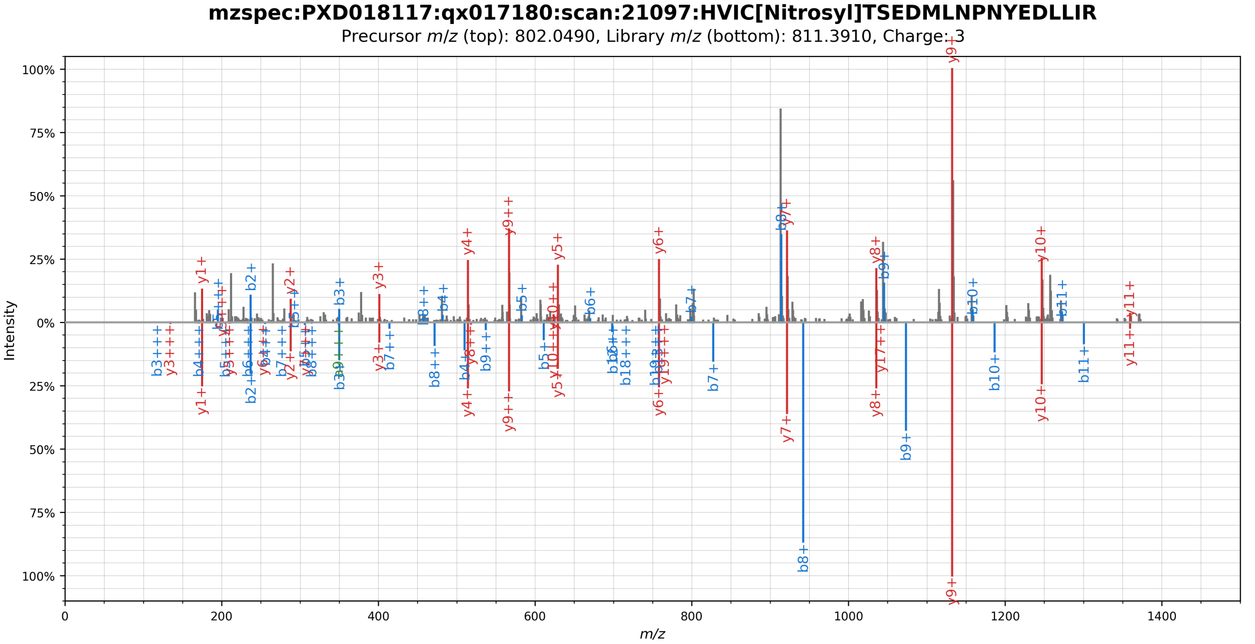


**Supplementary Figure 35.** Mirror plot showing the spectral library match (bottom) of a S-nitrosylated peptide originating from the SARS-CoV-2 Nsp5(C145A) protein (top). The S-nitrosylation was localized to residue C45 by visual inspection.

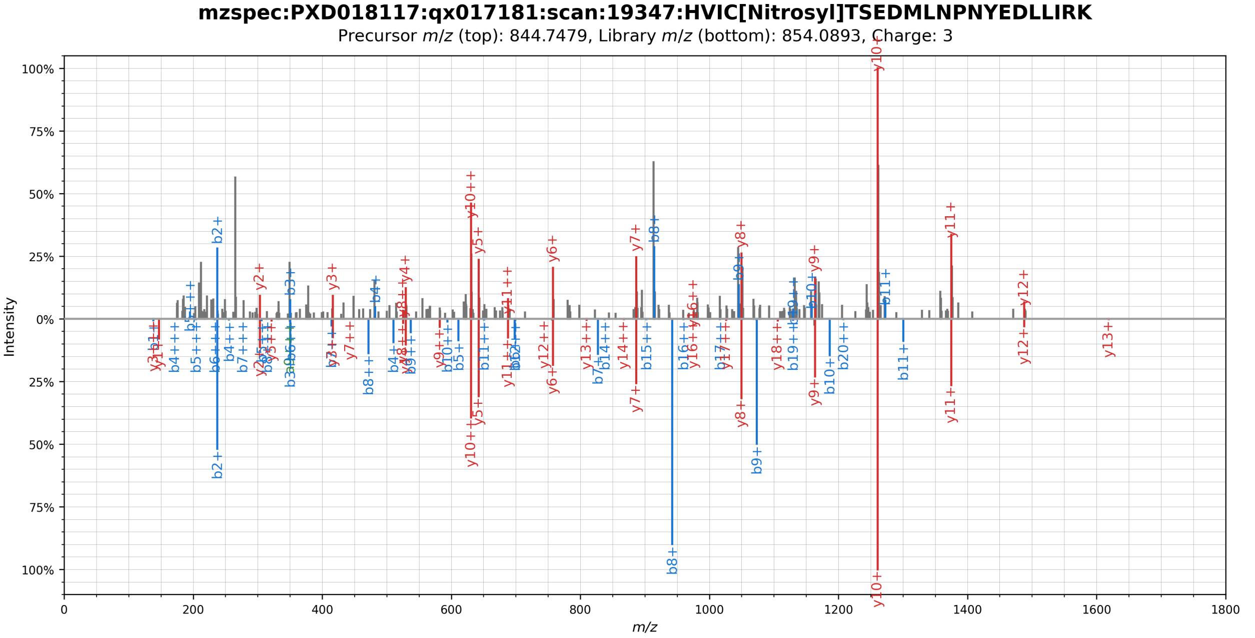


**Supplementary Figure 36.** Mirror plot showing the spectral library match (bottom) of a S-nitrosylated peptide originating from the SARS-CoV-2 Nsp5(C145A) protein (top). The S-nitrosylation was localized to residue C45 by visual inspection.

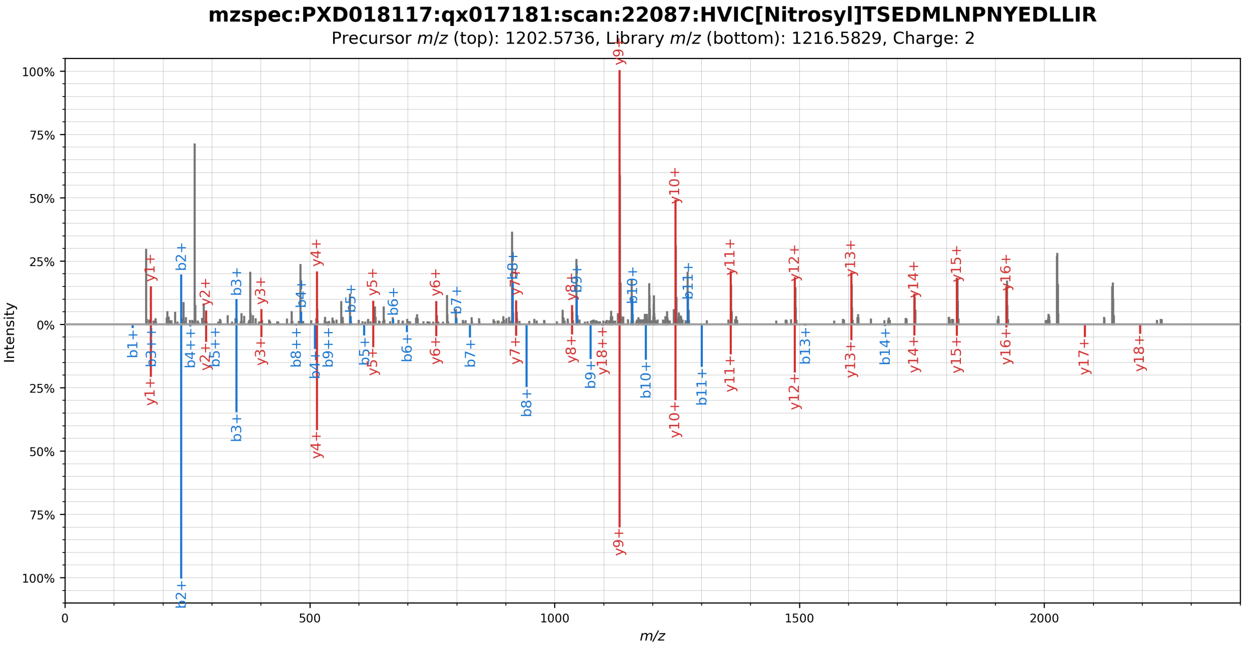


**Supplementary Figure 37.** Mirror plot showing the spectral library match (bottom) of a S-nitrosylated peptide originating from the SARS-CoV-2 Nsp5(C145A) protein (top). The S-nitrosylation was localized to residue C45 by visual inspection.

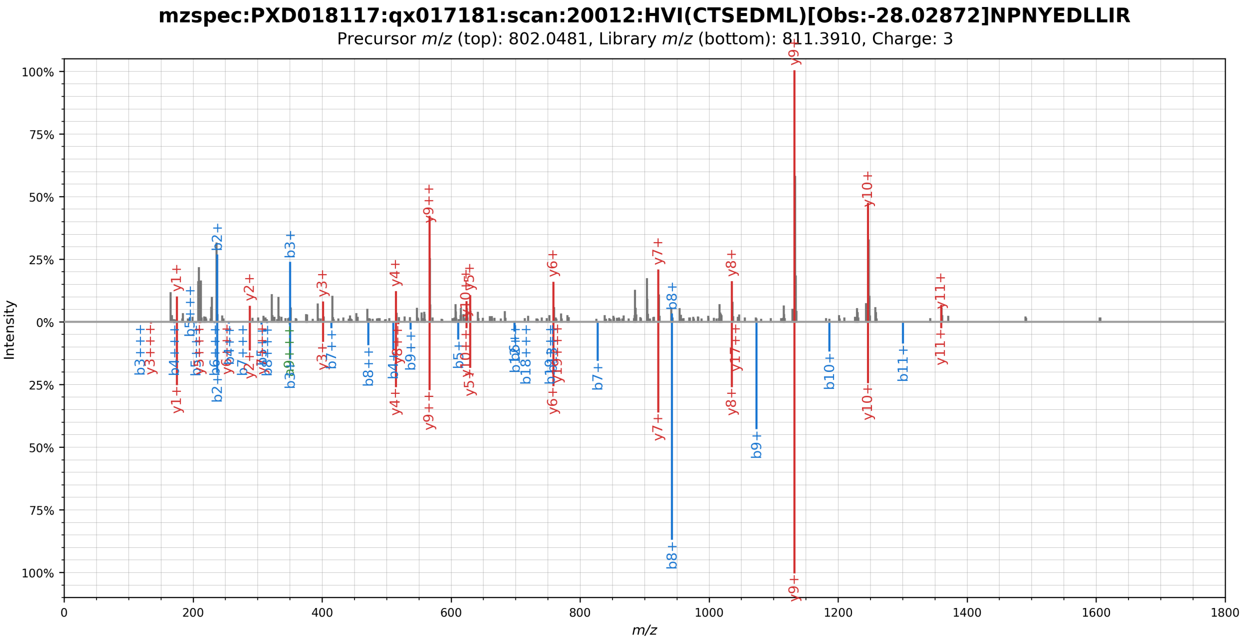


**Supplementary Figure 38.** Mirror plot showing the spectral library match (bottom) of a possibly S-nitrosylated peptide originating from the SARS-CoV-2 Nsp5(C145A) protein (top). The mass shift could not be localized by visual inspection.

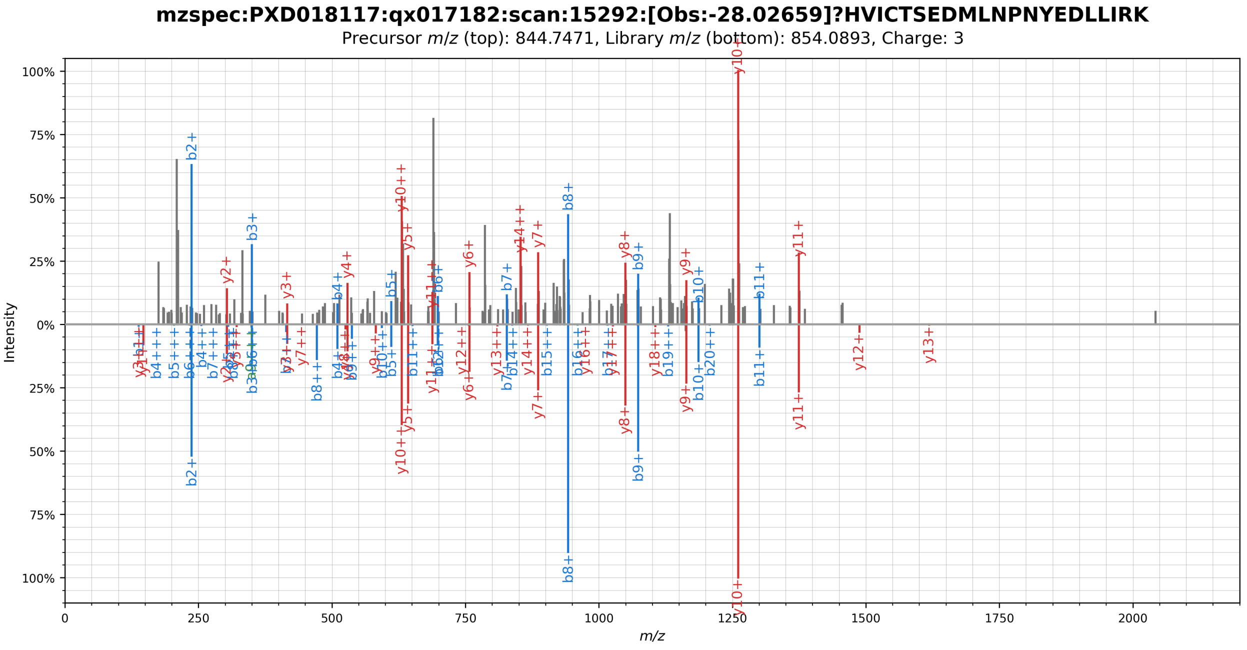


**Supplementary Figure 39.** Mirror plot showing the spectral library match (bottom) of a possibly S-nitrosylated peptide originating from the SARS-CoV-2 Nsp5(C145A) protein (top). The mass shift could not be localized by visual inspection.

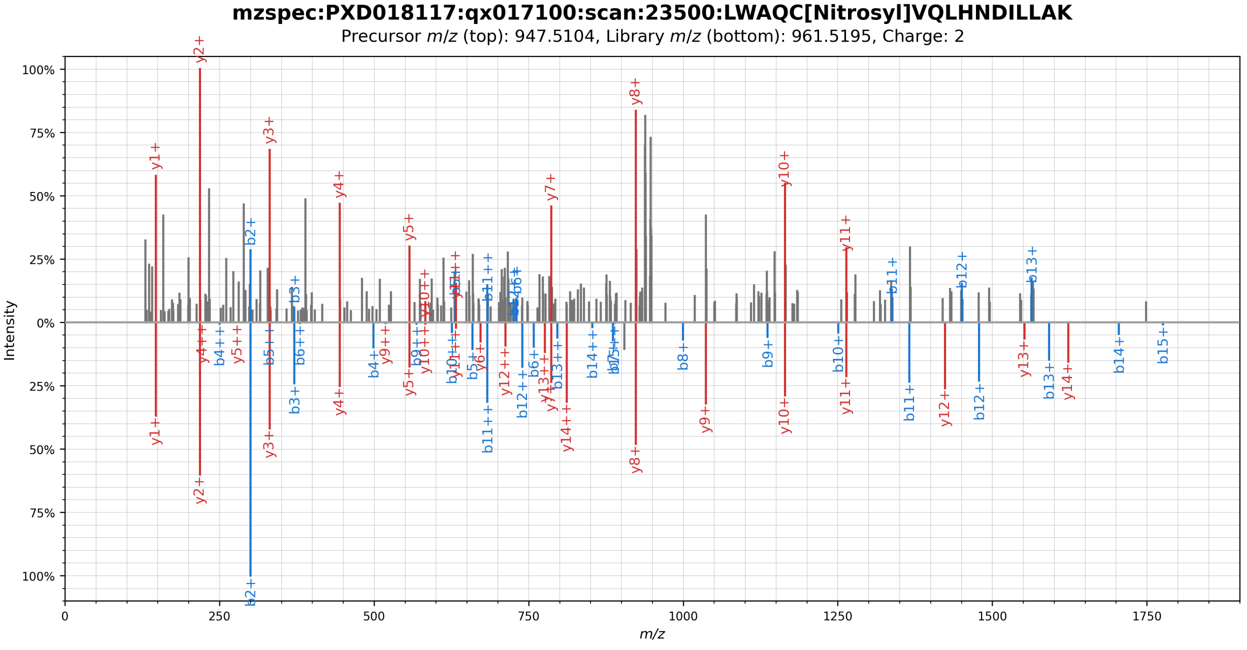


**Supplementary Figure 40.** Mirror plot showing the spectral library match (bottom) of a S-nitrosylated peptide originating from the SARS-CoV-2 Nsp7 protein (top). The S-nitrosylation was localized to residue C33 by visual inspection.

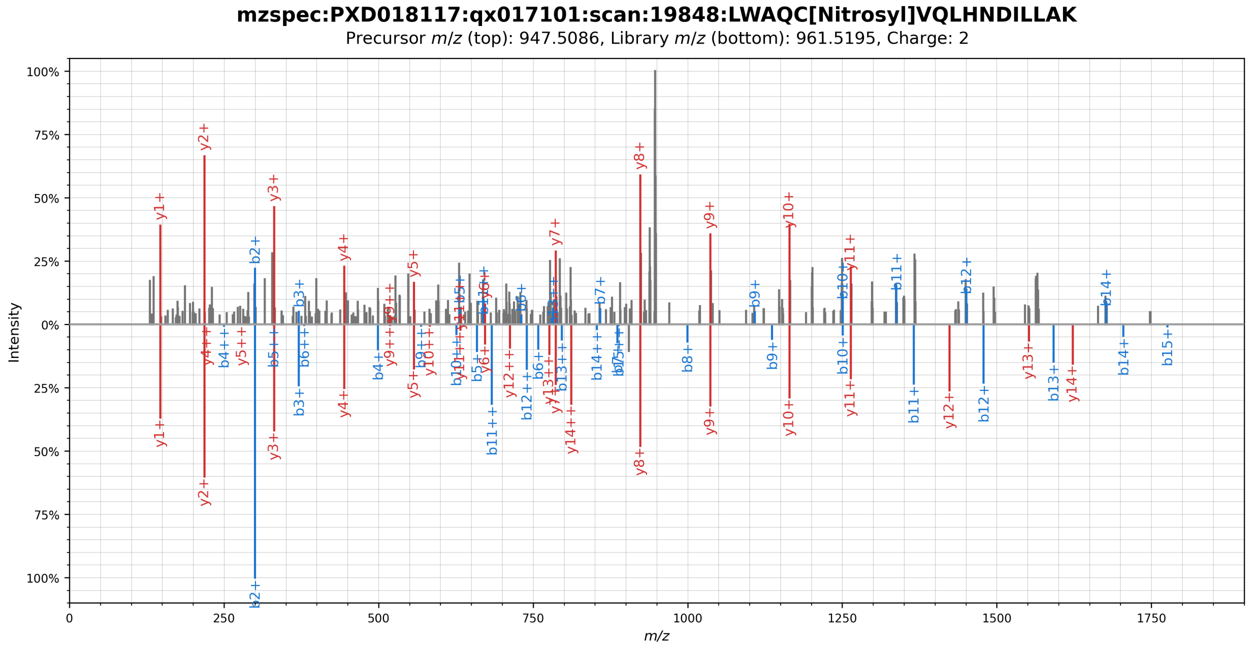


**Supplementary Figure 41.** Mirror plot showing the spectral library match (bottom) of a S-nitrosylated peptide originating from the SARS-CoV-2 Nsp7 protein (top). The S-nitrosylation was localized to residue C33 by visual inspection.

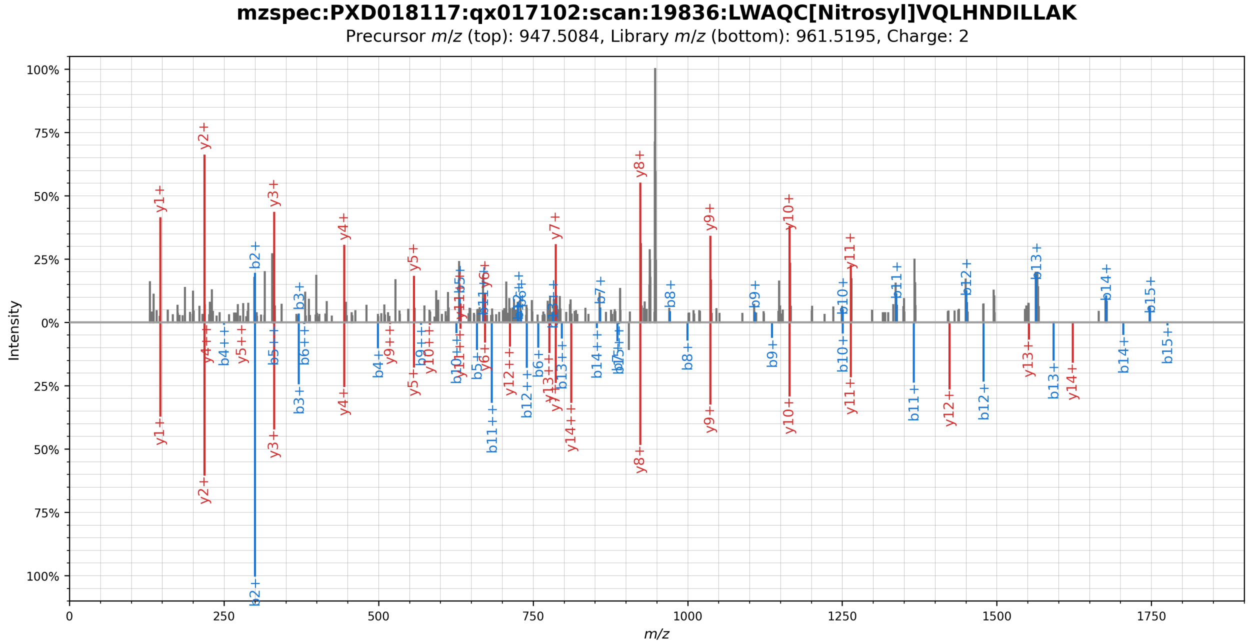


**Supplementary Figure 42.** Mirror plot showing the spectral library match (bottom) of a S-nitrosylated peptide originating from the SARS-CoV-2 Nsp7 protein (top). The S-nitrosylation was localized to residue C33 by visual inspection.

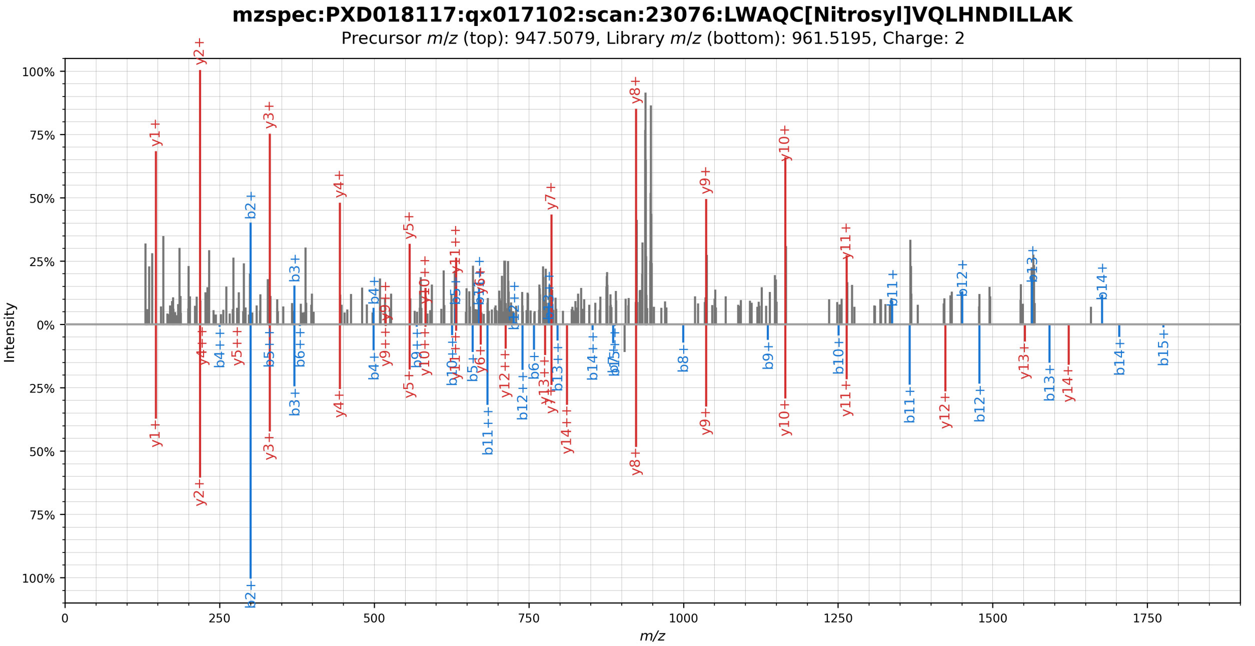


**Supplementary Figure 43.** Mirror plot showing the spectral library match (bottom) of a S-nitrosylated peptide originating from the SARS-CoV-2 Nsp7 protein (top). The S-nitrosylation was localized to residue C33 by visual inspection.

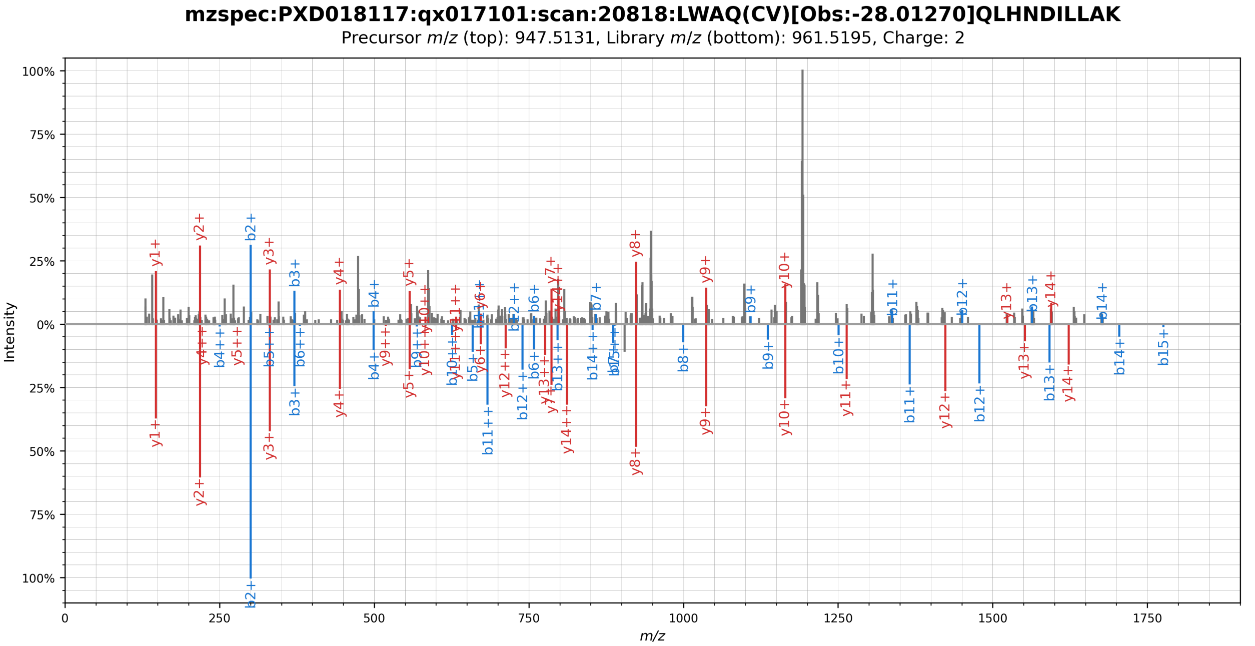


**Supplementary Figure 44.** Mirror plot showing the spectral library match (bottom) of a possibly S-nitrosylated peptide originating from the SARS-CoV-2 Nsp7 protein (top). The mass was localized to residue C33/V34 by visual inspection.

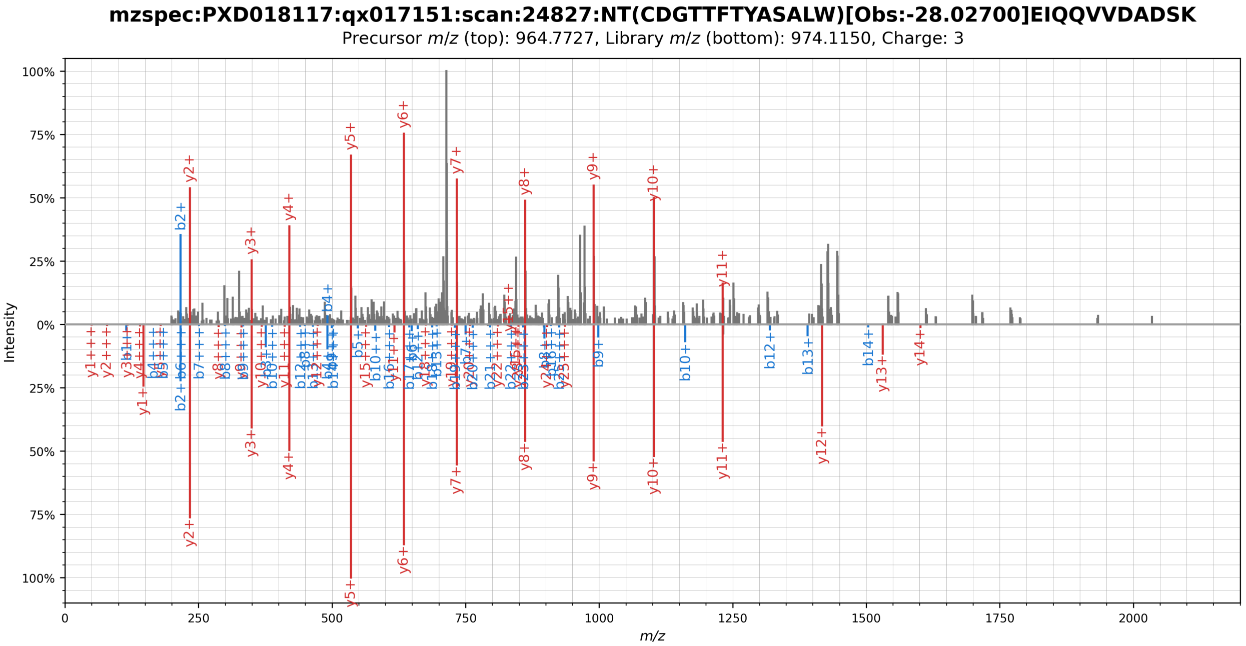


**Supplementary Figure 45.** Mirror plot showing the spectral library match (bottom) of a possibly S-nitrosylated peptide originating from the SARS-CoV-2 Nsp8 protein (top). The mass shift could not be localized by visual inspection.

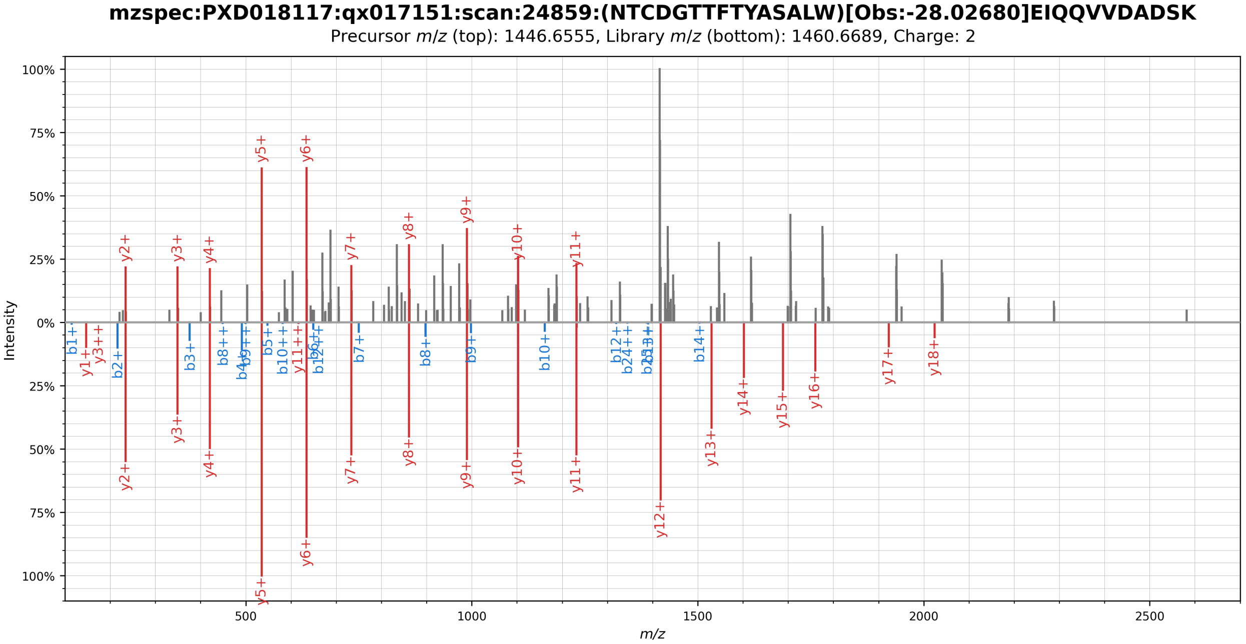


**Supplementary Figure 46.** Mirror plot showing the spectral library match (bottom) of a possibly S-nitrosylated peptide originating from the SARS-CoV-2 Nsp8 protein (top). The mass shift could not be localized by visual inspection.

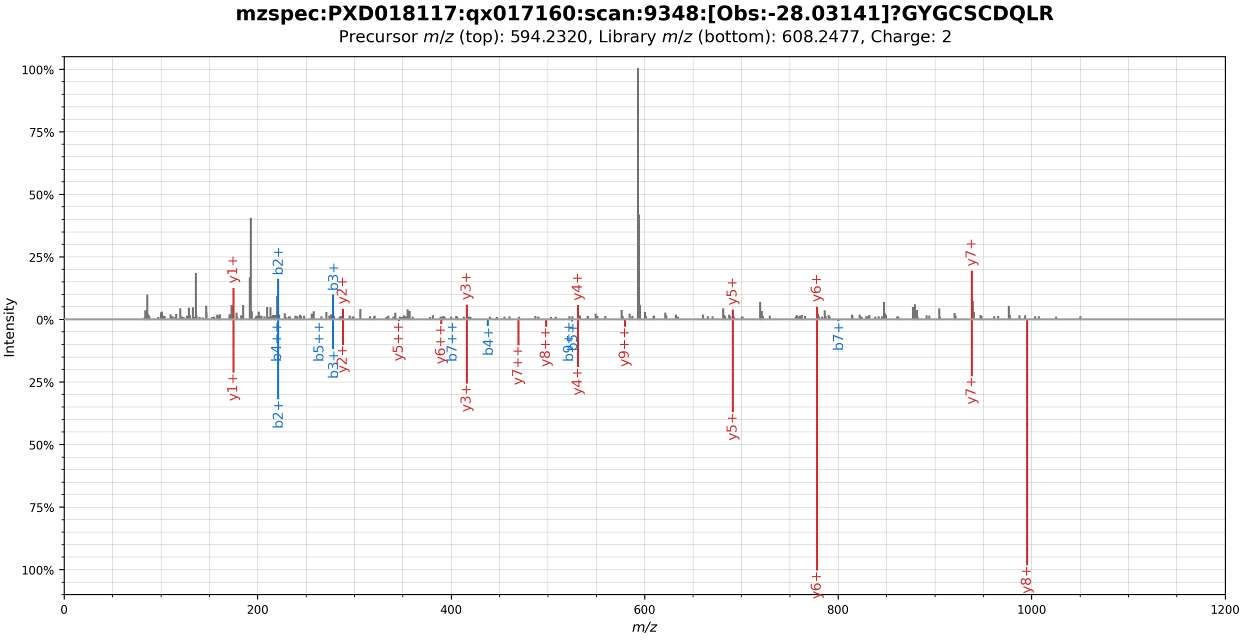


**Supplementary Figure 47.** Mirror plot showing the spectral library match (bottom) of a possibly S-nitrosylated peptide originating from the SARS-CoV-2 Nsp10 protein (top). The mass shift could not be localized by visual inspection.

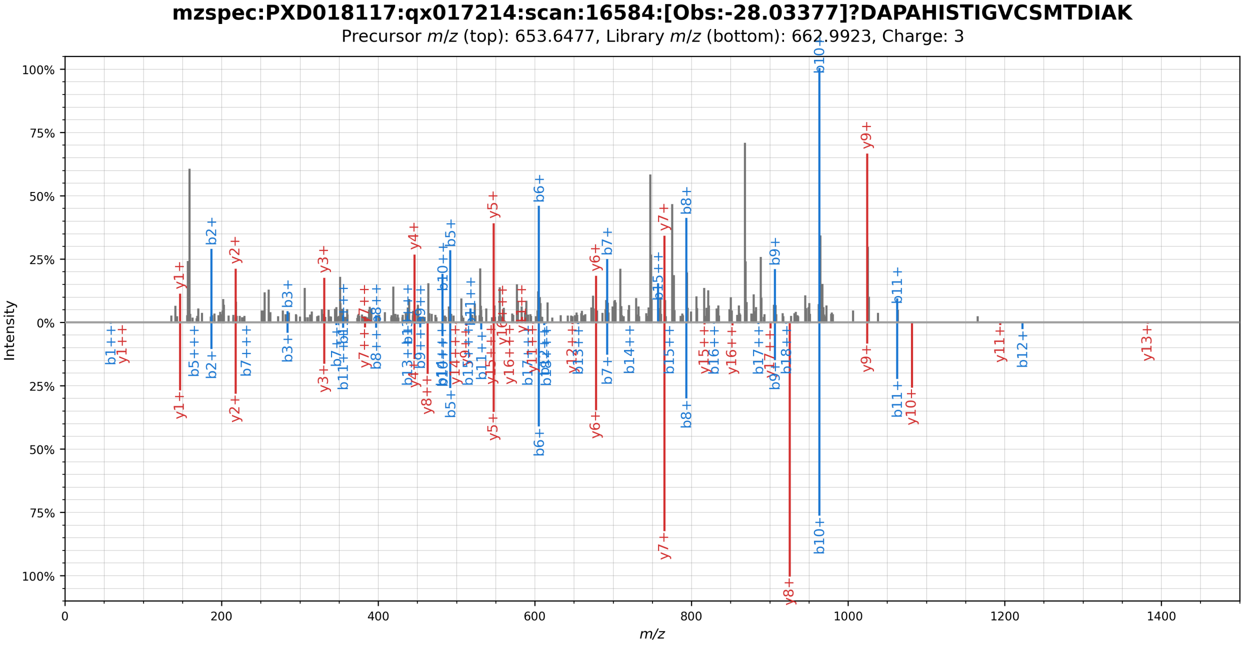


**Supplementary Figure 48.** Mirror plot showing the spectral library match (bottom) of a S-nitrosylated peptide originating from the SARS-CoV-2 Nsp15 protein (top). The S-nitrosylation was localized to residue C103 by visual inspection.

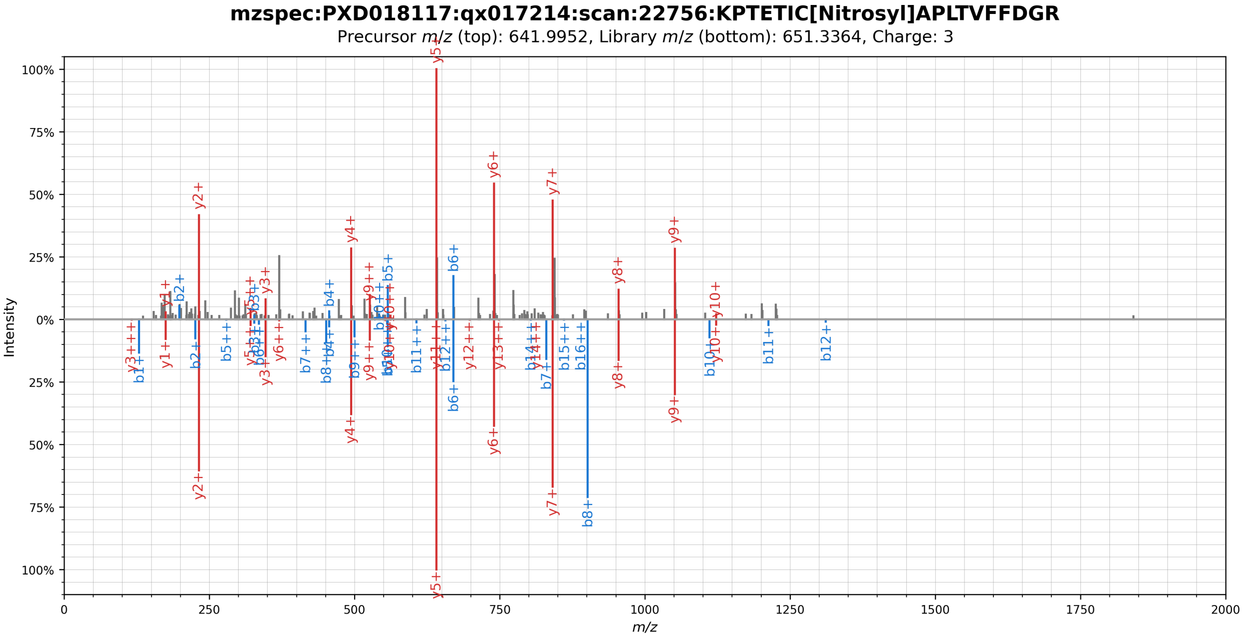


**Supplementary Figure 49.** Mirror plot showing the spectral library match (bottom) of a S-nitrosylated peptide originating from the SARS-CoV-2 Nsp15 protein (top). The S-nitrosylation was localized to residue C117 by visual inspection.

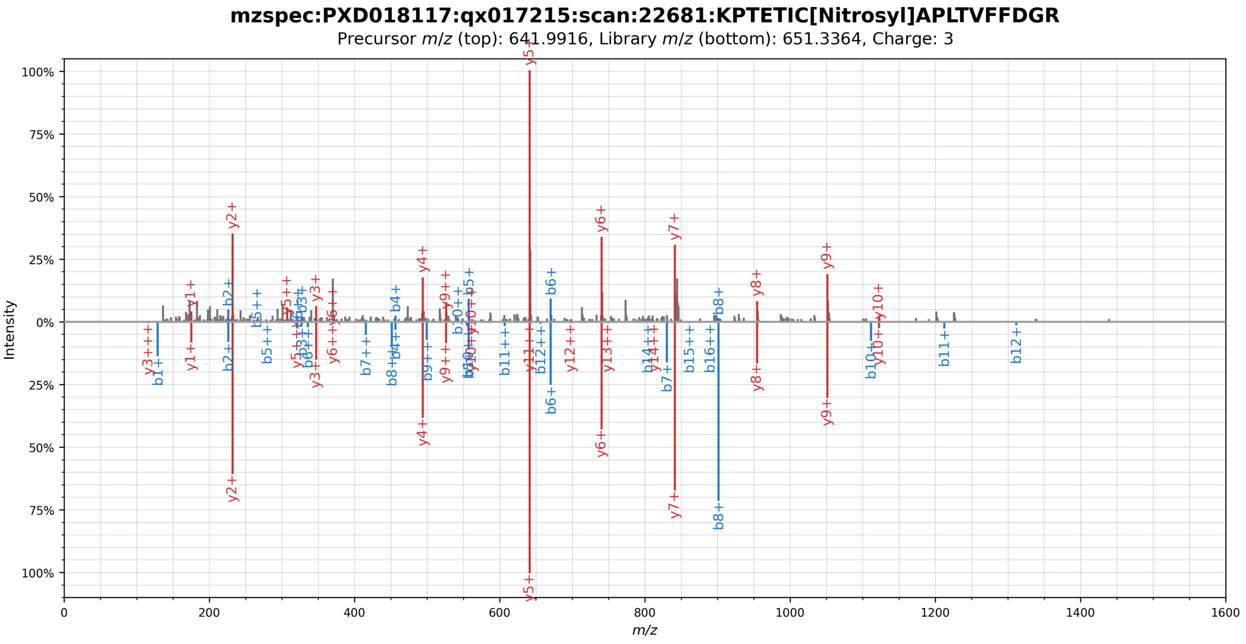


**Supplementary Figure 50.** Mirror plot showing the spectral library match (bottom) of a S-nitrosylated peptide originating from the SARS-CoV-2 Nsp15 protein (top). The S-nitrosylation was localized to residue C117 by visual inspection.

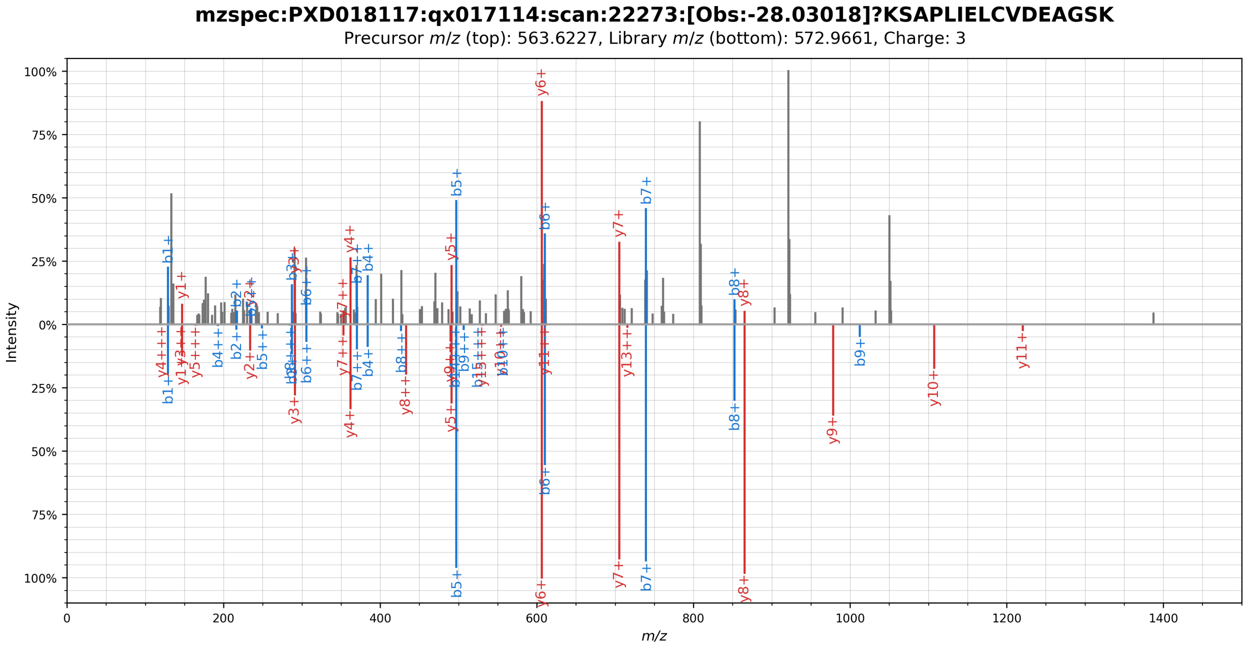


**Supplementary Figure 51.** Mirror plot showing the spectral library match (bottom) of a possibly S-nitrosylated peptide originating from the SARS-CoV-2 Orf8 protein (top). The mass shift could not be localized by visual inspection.

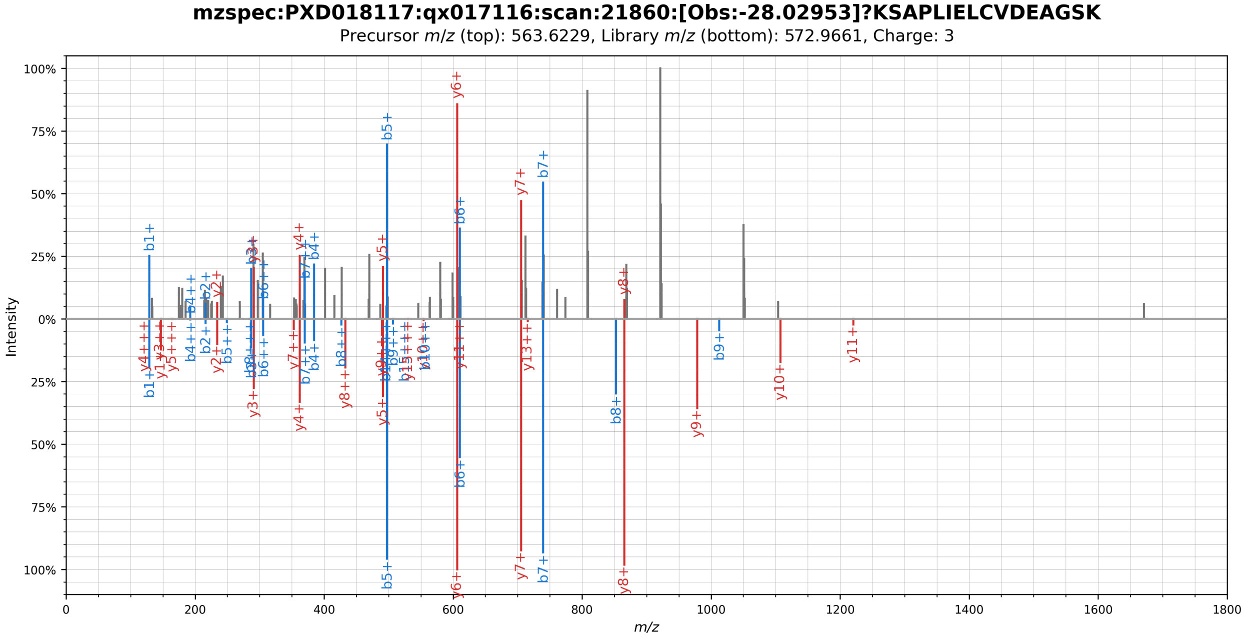


**Supplementary Figure 52.** Mirror plot showing the spectral library match (bottom) of a possibly S-nitrosylated peptide originating from the SARS-CoV-2 Orf8 protein (top). The mass shift could not be localized by visual inspection.
